# A novel subset of follicular helper-like MAIT cells has capacity for B cell help and antibody production in the mucosa

**DOI:** 10.1101/2020.10.05.326488

**Authors:** Owen Jensen, Shubhanshi Trivedi, Jeremy D. Meier, Keke Fairfax, J. Scott Hale, Daniel T. Leung

## Abstract

Mucosal-associated invariant T (MAIT) cells are innate-like T lymphocytes that aid in protection against bacterial pathogens at mucosal surfaces via release of inflammatory cytokines and cytotoxic molecules. Recent evidence suggests MAIT cells are capable of providing B cell help. In this study, we describe a previously unreported population of CXCR5^+^ T follicular helper (Tfh)-like MAIT cells, MAITfh, that have the capacity to provide B cell help within mucosal lymphoid organs. MAITfh cells are preferentially located near germinal centers in human tonsils and express the classical Tfh-associated transcription factor, B-cell lymphoma 6 (BCL-6), co-stimulatory markers, inducible T cell costimulatory (ICOS) and programmed death receptor 1 (PD-1), and cytokines, interleukin (IL)-21. Furthermore, we demonstrate the ability of MAIT cells to provide B cell help *in vivo* following mucosal challenge with *Vibrio cholerae*. Specifically, we show that adoptive transfer of MAIT cells into *αβ* T cell-deficient mice promoted B cell differentiation and increased serum *V. cholerae*-specific IgA responses. Our data demonstrate the capacity of MAIT cells to participate in adaptive immune responses, and suggest that MAIT cells may be potential targets for mucosal vaccines.

**One Sentence Summary:** We identified and characterized a novel subset of T follicular helper-like MAIT (MAITfh) cells that has the capacity to provide B cell help, and show the sufficiency of MAIT cells to promote production of pathogen-specific IgA antibodies and B cell differentiation in mucosal challenge.

## Introduction

Mucosal surfaces are in constant contact with both commensal and pathogenic microbes. One of the primary means of protection against invading pathogens is the production of secretory IgA and IgM by plasmablasts (PB) and plasma cells (PC) (*1*). In the lamina propria of the gut and lung, T independent low affinity IgA is produced against non-protein antigens by short-lived PBs. Conversely, T dependent high affinity antibodies against protein antigens are typically generated during germinal center reactions between T follicular helper (Tfh) and follicular B cells within local lymph nodes (LNs) or mucosal associated lymphoid tissues (MALT) (*2*). T follicular helper (Tfh) cells migrate to B cell follicles via upregulation of CXCR5 and downregulation of CCR7 (*3*). Within GCs, Tfh cells highly express the lineage-defining transcription factor, BCL6, and co-stimulatory molecules, PD1, ICOS, and CD40L (*4*). PD1 is important for Tfh GC positioning and function (*5*), while ICOS (*6*) and CD40L (*7*) engagement with B cells are essential for Tfh activation and GC formation (*4*). Tfh also produce cytokines, such as IL-21, that promote GC B cell responses (*8*).

Although Tfh cells are the primary drivers of T dependent GC responses, innate T cells, including invariant Natural Killer T (iNKT) cells and *γδ* T cells, also have Tfh like subsets capable of B cell help. In particular, iNKT cells have been well established in mice to provide both cognate and non-cognate help to B cells (*9–18*). Murine iNKTfh cells engage in cognate help leading to germinal center formation, and antibody class switching and production (*12, 14, 18*). iNKTfh cells have also been shown to promote non-cognate B cell help by licensing dendritic cells to recruit and activate Tfh cells (*10, 11*). CXCR5^+^ *γδ* T cells can promote antibody production *in vitro* (*19, 20*), and promote Tfh differentiation *in vivo* (*21*). More recent evidence suggests that another type of innate-like lymphocyte, Mucosal-associated invariant T (MAIT) cells, are capable of B cell help (*22–26*).

MAIT cells are innate-like *αβ* T cells defined by the expression of an invariant *α* chain, generally V*α*7.2 linked to J*α*33, 12 or 20 in humans and V*α*19 linked to J*α*33 in mice, and a limited array of TCR*β* chains (*27–29*). MAIT cells are highly enriched in human blood, liver and mucosa and are appreciated for their rapid ability to respond to microbial Vitamin B metabolites presented on the MHC class I related protein, MR1, or cytokine stimulation (*30–33*). Upon stimulation, MAIT cells produce pro-inflammatory cytokines including IFN*γ*, TNF*α*, and IL-17A and cytotoxic molecules including Granzyme B and Perforin (*34–36*). Furthermore, using MAIT deficient mice (MR1^-/-^), several studies have exemplified the importance of MAIT cells in immunity against mucosal bacterial pathogens (*37–40*).

Recent evidence suggests MAIT cells play a role in adaptive immune responses through B cell help. Analysis of human peripheral blood MAIT cells and serum antibody responses following *Vibrio cholerae* infection (*22*) and *Shigella* vaccination (*35*), revealed strong associations between MAIT frequency and activation with polysaccharide-specific IgA and IgG responses, but not with protein antibody responses (*22*). We have recently shown that human blood MAIT cells have the capacity to induce antibody production and B cell differentiation *in vitro*, and can secrete the B cell help cytokines following stimulation (*23*). Analysis of pleural effusions from tuberculosis patients revealed a population of PD1^High^ MAIT cells secreting key B cell help cytokines (*24*). Furthermore, two recent animal studies demonstrate the importance of MAIT cells in B cell help in murine autoimmunity (*25*) and mucosal vaccine immunity in non-human primates (*26*).

Our aims for this study were two-fold. We first wanted to determine whether a specific subset of MAIT cells are responsible for the B cell help phenotype. Our second aim was to determine *in vivo* if MAIT cells were sufficient to induce antibody production and humoral immune protection in the context of mucosal challenge. We found that like other innate-like T cells, MAIT cells have a Tfh like subset enriched within mucosal lymphoid organs. This MAITfh population expresses classical Tfh co-stimulatory markers, transcription factors, and cytokines, and is localized near B cell follicles. We further show, in a murine model, that adoptively transferred MAIT cells are capable of generating microbe-specific antibody responses against a mucosal bacterial pathogen in the absence of other *αβ* T cells.

Additionally, we find that in the context of mucosal challenge, MAIT cells promote increased production of microbe-specific IgA antibodies and mucosal B cell differentiation. These results suggest that MAIT cells have the capacity to enhance mucosal antibody mediated immunity, and thus may be a target in future mucosal vaccine development.

## Results

### CXCR5^+^ MAIT cells are increased in tonsils and express higher levels of Tfh co-stimulatory markers compared to peripheral blood

With recent evidence highlighting the ability of MAIT cells to provide B cell help, and studies showing a modest population of MAIT cells within human and mouse mucosal lymphoid tissues (*39, 41, 42*), we aimed to determine if MAIT cells had a Tfh-like subset capable of B cell help. To investigate this, we used flow cytometry to analyze MAIT cell expression of classical Tfh markers in human peripheral blood mononuclear cells (PBMCs) and tonsils. We obtained PBMC’s from adult blood donors, and tonsils from children, ages 2-16, undergoing tonsillectomy for recurrent tonsillitis or tonsillar hyperplasia. MAIT cells were defined as CD3^+^ V*α*7.2^+^ MR1-5-OPRU Tetramer^+^ cells and gated based on a PE conjugated MR1-6FP Tetramer negative control (fig S1A, S1B and Fig. 1A). Median MAIT frequency among total CD3^+^ cells in tonsils was 0.23% (interquartile range (IQR) = 0.14%, 0.29%) compared to 1.03% (IQR = 0.62%, 1.35%) among PBMCs (Fig. 1B). We found that a higher percentage (Median=18.1%, IQR=9.2%, 28.6%) of tonsil MAIT cells were CXCR5^+^ compared to PBMC MAIT cells (Median=1.74%, IQR=0.37%, 2.16%, p<0.0001), although there was considerable variability among tonsil MAIT cells (Fig. 1C and fig. S1C). To further confirm the MAIT cell phenotype, we measured expression of two C type lectin receptors, CD161, a common delineator of MAIT cells (*30*), and CD69, a marker of T cell tissue residency (*43*). We found that significantly fewer CXCR5+ MAIT cells expressed CD161 in both PBMC (p<0.01) and tonsils (p<0.001) than did CXCR5-MAIT cells (fig. S2D and F). The loss of CD161 expression has been demonstrated in vitro following TCR stimulation, suggesting CXCR5+ MAIT cells may be previously activated (*44*). Despite being MR1-Tetramer+ and TRAV1-2+, we wanted to confirm that the CD161-fraction were indeed MAIT cells and therefore utilized paired TCR*αβ*single-cell Illumina sequencing as previously described (*45*). We sorted and single-cell sequenced CD161+ and CD161-MAIT cells (identified as live CD3^+^ V*α*7.2^+^ MR1-Tetramer^+^ cells) from four tonsil samples and analyzed *TRAJ* expression in *TRAV1-2* expressing wells. We found no statistically significant differences in *TRAJ* expression frequency between CD161+ and CD161-MAIT cells, (fig. S3A) with the majority expressing *TRAJ-33* with average frequencies of 66.3% ± 17.5 and 65.5% ± 7.9, respectively. In concurrence with previous studies (*29, 46–48*), *TRAJ-34* and *TRAJ-12* were also highly expressed in both groups highlighting that these are bona fide MAIT cells. Furthermore, we found no statistically significant differences in the percentages of CXCR5+ and CXCR5-MAIT cells that expressed CD69 in PBMC and tonsils (fig. S2C and E), though both CXCR5+ and CXCR5-MAIT cells in the tonsils had higher cell surface levels CD69 expression, indicating a tissue resident memory (Trm) phenotype. We next wanted to determine if there were differences in Tfh co-stimulatory marker expression between CXCR5+ and CXCR5-MAIT populations. Significantly more CXCR5+ MAIT cells in tonsils expressed ICOS (p<0.0001) (Fig. 1D and fig S2B) than did CXCR5-MAIT cells; moreover, significantly more CXCR5+ MAIT cells in both tonsils and PMBC’s expressed PD1 (Fig. 1E and fig S2A) than did CXCR5-MAIT cells in the same tissue. Additionally, significantly more CXCR5+ MAIT cells in tonsils expressed PD1 (p<0.0001) and ICOS (p<0.0001) than did PBMC CXCR5+ MAIT cells. Notably, ICOS and PD1 frequency and fluorescence intensity among tonsil CXCR5^+^ MAIT cells are similar to that seen in tonsil CD4^+^ CXCR5^+^ (Tfh) cells (Fig.1D and 1E, and fig. S2A and S2B). In order to confirm that the differences between PBMC and tonsil MAIT cells are not age dependent, we analyzed MAIT cell CXCR5, ICOS and PD1 expression in 5 paired PBMC and tonsil samples from pediatric donors ages 3-15 (fig. S4). Similar to adult PBMC samples we found increased total MAIT frequency and decreased CXCR5 expression in PMBC compared to tonsil samples (fig. S4B-C). Similar trends in ICOS and PD1 expression were found in the pediatric PBMC MAIT and MAITfh groups compared to the adult PBMC donors supporting the use of adult PBMC samples in the following figures where low pediatric blood sample volumes would limit analysis (Fig. 1D-E and fig. S4D-E). Taken together, a high proportion of MAIT cells in tonsils have a CXCR5^+^ phenotype with Tfh-like co-stimulatory markers.

**Fig. 1.**
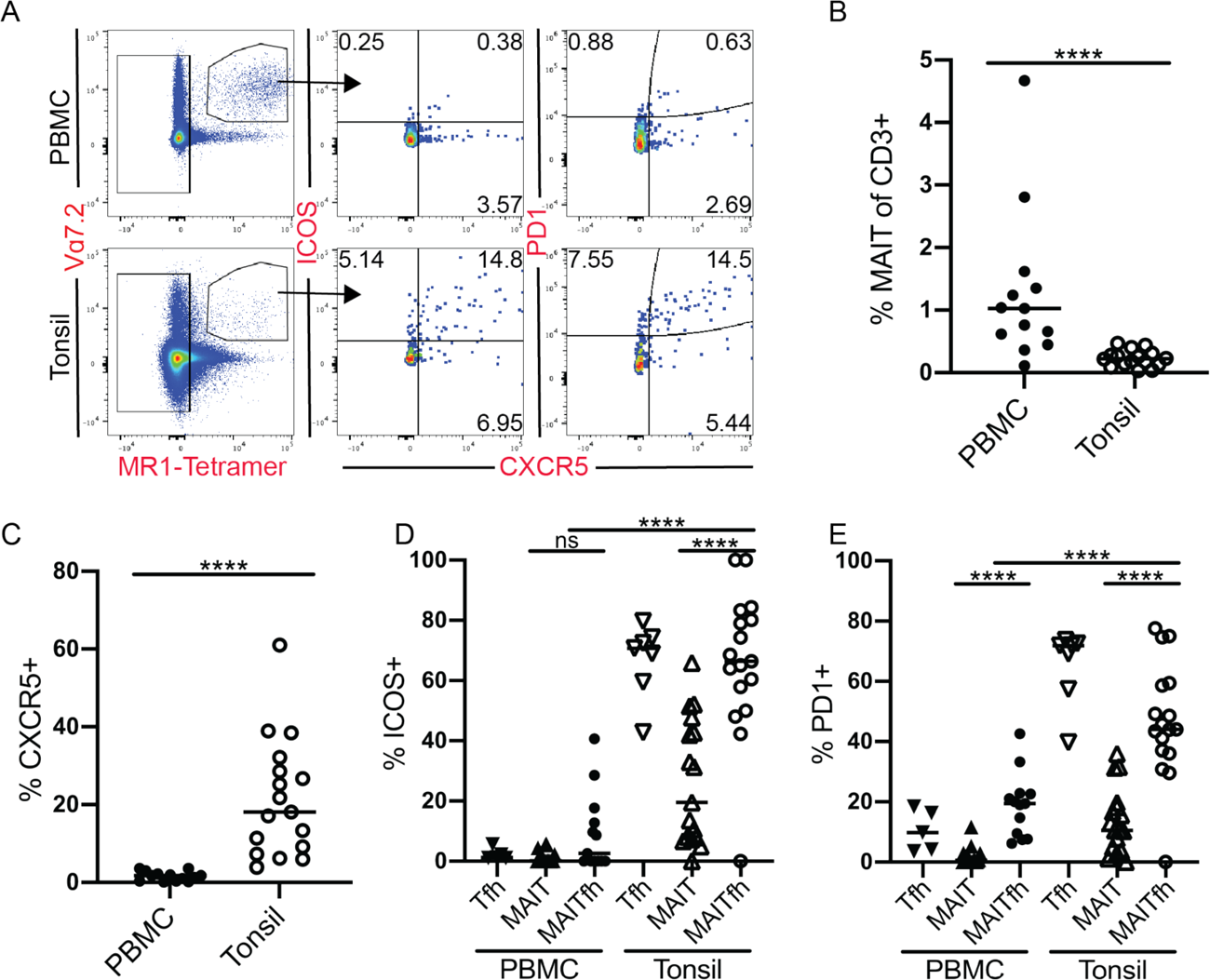
Increased expression of Tfh co-stimulatory molecules in tonsil MAIT cells. (**A**) Representative FACS plots of unstimulated CD3^+^ PBMC (top) and tonsil (bottom) cells with MAIT cell (gated in left panel) co-expression of CXCR5^+^ with ICOS (middle) and PD1 (right). (**B**) Quantification of MAIT cell frequency as percentage of CD3^+^. (**C**) Frequency of CXCR5^+^ MAIT cells. Frequency of (**D**) ICOS and (**E**) PD1 MAIT (CXCR5^-^), MAITfh (CXCR5^+^) and Tfh cells. Data are represented as Median from 2 pooled independent experiments. n *≥* 13. *\*p* < 0.05, *\*\*p*<0.01, *\*\*\*p*<0.001, *\*\*\*\*p*<0.0001 by two-tailed Mann-Whitney *U* test

### Transcriptional analyses of CXCR5^+^ MAIT cells reveals expression of Tfh associated cytokines and transcription factors

We have previously demonstrated that peripheral blood MAIT cells can secrete B cell help cytokines IL-6, IL-10 and low levels of IL-21 *in vitro* (*23*). MAIT cells are also potent producers of IFN*γ* (*49*), which is known to have both inhibitory and activating roles in B cell development, proliferation, and antibody responses (*50*). In order to investigate the transcription factor and cytokine expression of CXCR5^+^ and CXCR5^-^ MAIT cells, we examined transcript levels of fluorescent associated cell sorted (FACS) MAIT and non-MAIT CD3^+^ T cell populations from tonsil and blood. We refer to the sorted populations, as CD8^+^ (CD8^+^ CXCR5^-^), Tfh (CD4^+^ CXCR5^+^), MAIT (MR1-Tet^+^, V*α*7.2^+^, CXCR5^-^), and MAITfh (MR1-Tet^+^, V*α*7.2^+^, CXCR5^+^) (fig. S5A). To confirm our flow analysis, we assayed for *CXCR5* expression. We found significant increases in *CXCR5* relative transcript levels in the MAITfh compared to the MAIT groups in both PBMC’s (p<0.01) and tonsils (p<0.001) (Fig. 2A). Though notably, the MAITfh group had lower CXCR5 expression than the control Tfh group in both tissues. We next assayed for the Tfh lineage-defining transcription factor (TF), *BCL6*. The MAITfh population within PBMCs had higher *BCL6* expression compared to MAIT (p<0.05) and Tfh (p<0.01) cells. Within tonsils, MAITfh cells had higher *BCL6* expression compared to Tfh (p<0.01) and CXCR5-MAIT (p<0.08) populations (Fig. 2B). In comparison, analysis of canonical T cell associated TFs *RORC* (Th17) (p<0.05), and *TBX21* (Th1) (p<0.05), revealed higher expression in the tonsil MAIT over the MAITfh population (Fig. 2C and D).

**Fig. 2.**
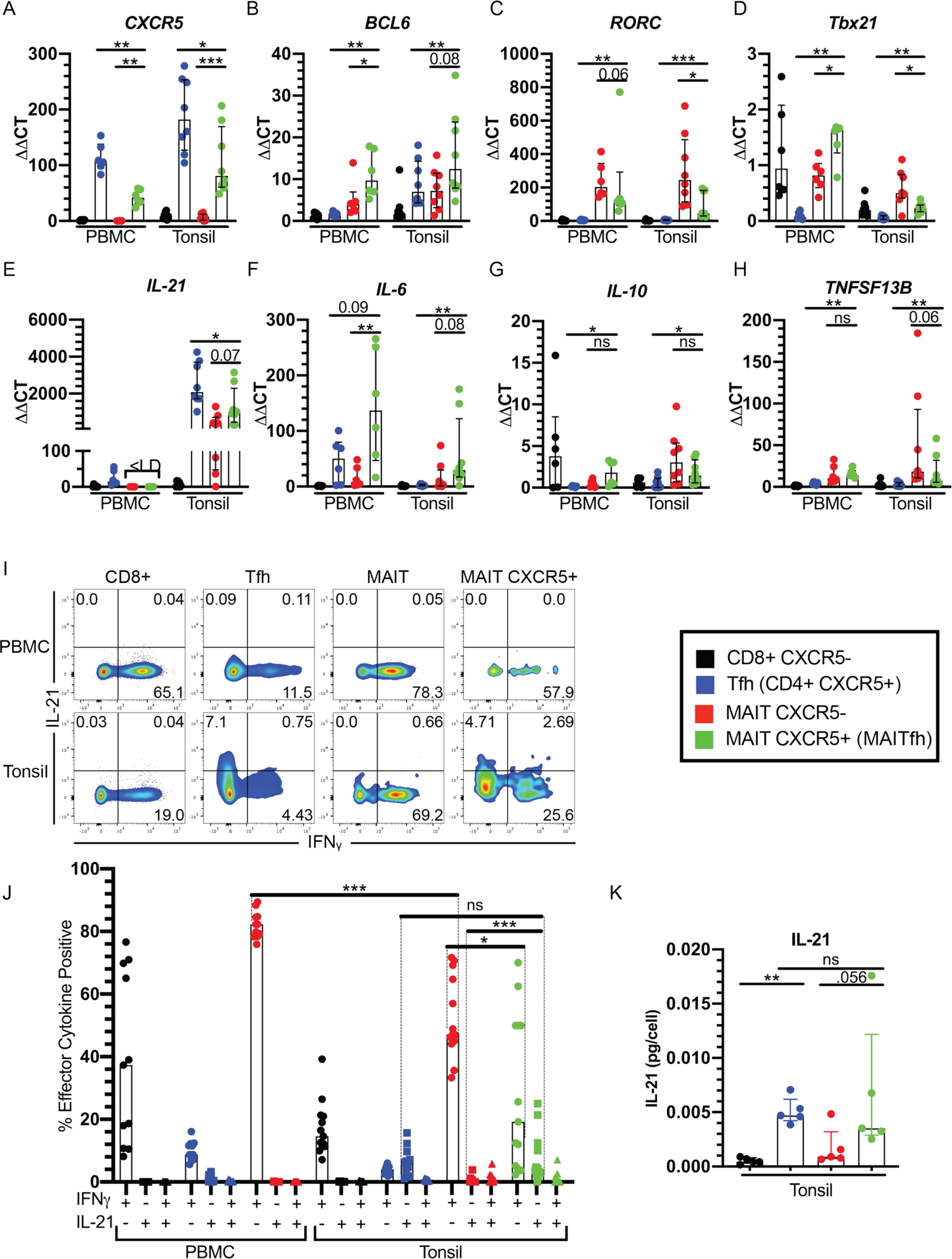
CXCR5^+^ MAIT cells highly express B cell help cytokine IL-21. (A) *CXCR5*, (B) *BCL6*, (C) *RORC*, (**D**) *TBX21*, (**E**) *IL-21*, (**F**) *IL-6*, (**G**) *IL-10*, and (**H**) *TNFSF13B* qPCR data of unstimulated FAC sorted PBMC and tonsil T cell populations. Data are represented as *ΔΔ*Ct relative to an internal housekeeping gene, *β*-Actin, and the PBMC-CD8^+^ CXCR5^-^ population. Data are Median with IQR from 2 independent experiments. n=6-8. (**I**) Representative flow cytometry from PBMC and tonsil T cell populations of IL-21 and IFN*γ* co-expression following 6 h stimulation with PMA/Ionomycin and Brefeldin A for final 4h. (**J**) Frequency of T cell populations expressing IL-21, IFN*γ* or both. Data are Median with IQR from 2 pooled independent experiments. n*≥*11. (**K**) IL-21 ELISA following 42 h PMA/Ionomycin stimulation of FAC sorted mono-cultured T cell populations from tonsils described below. Data are Median with IQR, n=5. T cell populations described gated on live CD3+ cells are as follows: Black =CD8^+^ CXCR5^-^, Blue = Tfh (CD4^+^ CXCR5^+^), Red= MAIT (MR1-Tetramer^+^ Va7.2^+^) CXCR5^-^, Green= MAIT CXCR5^+^. Representative gating for populations found in Sup. Fig 3a. *p < 0.05, **p<0.01, ***p<0.001, ****p<0.0001 by two-tailed Mann-Whitney U test.

MAITfh groups had higher expression of *RORC* and *TBX21* compared to Tfh groups in PBMCs (p<0.01) and tonsils (p<0.01) (Fig. 2C and D). Given that conventional Tfh cells are known to co-express TFs associated with other helper T cell subsets based on lineage and environment (*51–53*), our data suggest that co-expression of non-follicular helper-associated TFs in the MAITfh population may suggest potential for plasticity in phenotype or differentiation from a Th1 or Th17-like subset.

We next assayed for cytokines known to play a role in B cell differentiation or antibody production. *IL-21* expression, although below the limit of detection (<LD) in PBMC MAIT and MAITfh groups, had a trending increase in the tonsil MAITfh compared to the MAIT groups (p<0.07) (Fig. 2E). Notably, although 2-5 fold lower than the tonsil Tfh group, both tonsil MAIT (median=361.3) and MAITfh (Median=977.9) groups highly expressed *IL-21* relative to the tonsil CD8^+^ CXCR5^-^ group (Median=6.0) (Fig. 2E). We found similar increases in *IL-6* expression in both PBMC and tonsil MAITfh compared to MAIT and Tfh groups (Fig. 2F). Interestingly, both *TNFSF13B (BAFF)* and *IL-10* expression were significantly higher in both MAIT and MAITfh groups within tonsils and PBMCs compared to Tfh cells in their respective compartments (Fig. 2G and H). Taken together, CXCR5^+^ MAIT (MAITfh) cells express Tfh-associated cytokines and transcription factors at levels similar or higher than Tfh cells in their respective compartments

### CXCR5^+^ MAIT cells highly express IL-21 compared to CXCR5^-^ MAIT cells

We next wanted to investigate the relationship between IL-21 and IFN*γ* expression among MAIT subsets as both cytokines are highly expressed and generally associated with a Tfh vs Th1 like phenotype. When we stimulated PBMCs with PMA/Ionomycin, we saw a clear dichotomy between IFN*γ* and IL-21 positive cells in all populations with very low frequency of double positive cells, and the vast majority being IFN*γ*^+^ (Fig. 2 I and J). MAITfh cells seemed to have a substantially lower frequency of IFN*γ*^+^ cells compared to conventional MAIT cells, though these data were limited by low PBMC-MAITfh cell numbers, potentially biasing results. This effect was also notable in tonsils (p<0.001), where IFN*γ* production in MAITfh cells was significantly reduced compared to that of MAIT cells (p<0.05), and IL21 production was significantly increased (p<0.001). Very few double-positive cells were recorded in either population. Furthermore, the tonsil MAITfh population had a similar frequency of IL-21^+^ cells compared to conventional Tfh cells following stimulation (Fig. 2I and J). IFN*γ*^+^ and IL21^+^ frequency were low in all unstimulated populations (fig. S6A-B). Conversely, when stimulated with a combination of 5-amino-6-D-ribitylaminouracil (5-A-RU) and methylglyoxal to simulate MR1-dependent MAIT activation through formation of the MAIT ligand, 5-OP-RU (*54*), we found minimal IL-21^+^ MAIT or MAITfh cells and statistically similar IFN*γ*^+^ frequencies (fig. S7A-B). This suggests that IL-21 production by MAIT cells may be MR1-independent, or dependent on alternative stimulation conditions. To confirm the IL-21 production capacity of MAIT cells and MAITfh cells within tonsils, we sorted MAIT (CD3^+^ V*α*7.2^+^ MR1-Tetramer^+^ CXCR5^-^), MAITfh (CD3^+^ V*α*7.2^+^ MR1-Tetramer^+^ CXCR5^+^), CD8^+^ CXCR5^-^ and GC Tfh (CD4^+^ CXCR5^+^ PD1^high^) populations, and stimulated mono-cultures of 17,000-40,000 cells with PMA/Ionomycin (*55*). To account for differences in cell number per well we normalized all data to pg IL-21 per cell. We found a trending increase in IL-21 production in the MAITfh over MAIT groups (p<0.056) and no statistical difference in IL-21 production between MAITfh and GC Tfh populations (Fig. 2K). Taken together, these data demonstrate the potent ability of the tonsil MAITfh population to produce B-cell help cytokines and dichotomy in phenotype between classical MAIT cells and MAITfh cells in both peripheral blood and tonsil tissue.

### PD1^+^ MAIT cells localize near germinal centers within tonsils

Having established that CXCR5^+^ MAIT cells exhibit a Tfh-like phenotype *in vitro*, we next sought to determine the *in vivo* spatial differences between CXCR5^-^ and CXCR5^+^ MAIT cells in relation to germinal centers (GCs) in tonsil cryosections. MAIT cells were defined by co-staining of TRAV1-2 and CD161 as described in Leng et al. 2019 (*56*). Anti-IgD was used to stain naïve B cells making up the follicular mantle, and potential IgD^+^ memory B cells in the marginal or superficial zone (*57*). Due to difficulties with co-staining MAIT cells and CXCR5, we used PD1 to mark MAITfh and non-MAIT GC Tfh cells (Fig. 3A). To confirm PD1 expression on Tfh cells, we stained tonsil cyrosections with anti-CD4, anti-CD3, anti-IgD and anti-PD1. We found high co-expression of CD3^+^ CD4^+^ cells and PD1 within tonsil GCs, while minimum PD1 expression was found in the CD3^+^ CD4^+^ cells outside of the follicular mantle (fig. S8A). These data suggest that GC Tfh cells can be marked by high PD1 expression. PD1 has also been used to mark GC light zone Tfh cells in human adenoid samples (*58*) and non-human primates (*59*). Furthermore, flow cytometry analysis revealed significantly higher expression of PD1 among CXCR5^+^ (Median=44.1%, IQR=37.1%, 49.2%) vs CXCR5^-^ (Median=10.5%, IQR=4.88%, 18.2%) MAIT cells (Fig. 1E). ImageJ was used to quantify and measure PD1^+/-^ MAIT cell distance to the edge of the nearest GC, outlined in Fig. 3A (dashed white line). A total of 349 MAIT cells from n=5 tonsils were analyzed with 32% ± 15.9 being PD1^+^. In comparison, flow results of unstimulated tonsil cells showed 38% ± 14.9 of MAIT cells to be PD1^+^, thus confirming the ability of the immunohistochemistry imaging and quantification to assess this cell population. PD1^+^ MAIT cells were generally located within or closely surrounding GC’s with a median distance of 66.8 μm from a GC edge compared to PD1^-^ MAIT cells with a median distance of 220.2 μm (Fig. 3B). 28.4% of PD1^+^ MAIT cells were located within GC’s and thus were recorded as 0 μm. PD1^High^ expressing MAIT cells tended to localize with other PD1^High^ cells within GC light zones. In contrast, PD1^low^ or PD1^-^ MAIT cells were often found outside of the IgD^high^ follicular mantle zone (Fig. 3A-yellow dashed line) and frequently in contact with IgD^low^ cells. Overall, combined with phenotypic analysis above, these data reveal PD1^+^ (and likely CXCR5^+^) MAIT cells to be Tfh-like in their proximity to germinal centers

**Fig. 3.**
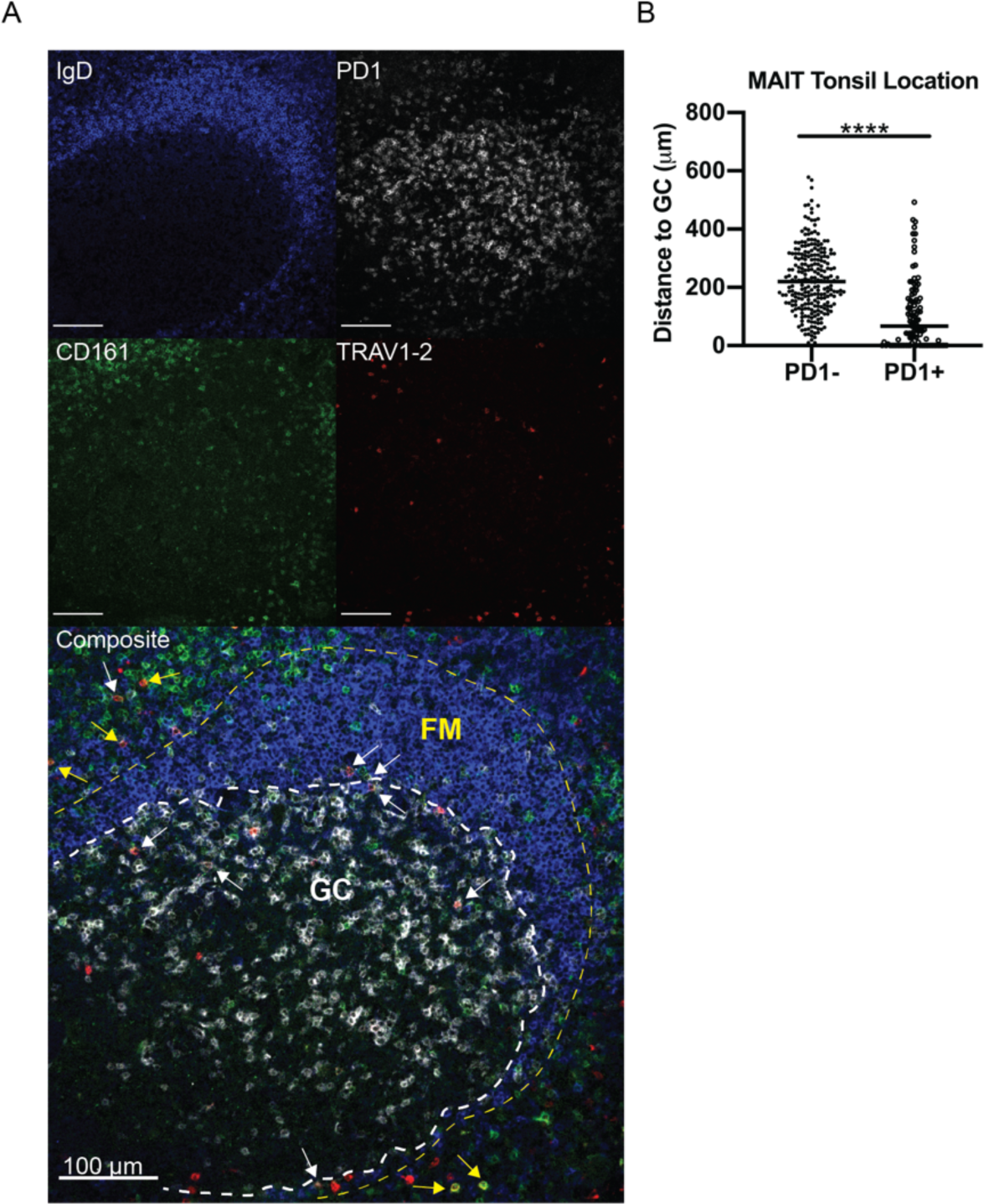
PD1^high^ MAIT cells preferentially locate near B cell follicles. (**A**) Location of MAIT cells within tonsils. Representative immunofluorescence images from tonsil cryosections stained with anti-IgD (blue), anti-PD1 (white), anti-CD161 (green), and anti-TRAV1-2 (red) and imaged on a Leica SP8 confocal microscope with 20X objective. Enlarged composite image below with MAIT cells (TRAV1-2^+^ CD161^+^) denoted with arrows. White arrows indicate PD1^+^ MAIT cells. Yellow arrows indicate PD1^-^ MAIT cells. White dashed line outlines germinal center (GC) highlighted by PD1^high^ cells. Yellow dashed line outlines follicular mantle (FM) highlighted by IgD^+^ naïve B cells. All scale bars represent 100 μm. (**B**) Quantification of MAIT cell distance to edge of nearest GC. Tiled images from tonsil donors were visually assessed for co-staining of CD161, TRAV1-2 and then PD1. Distance to nearest GC was then measured using ImageJ. n=359 MAIT cells were quantified (PD1^+^ = 103; PD1^-^ = 246) from n=5 tonsil donors. Data are represented as medians with dots representing single MAIT cells. *\*p* < 0.05, *\*\*p*<0.01, *\*\*\*p*<0.001, *\*\*\*\*p*<0.0001 by two-tailed Mann-Whitney *U* test.

### MAIT cells repopulate mucosal and lymphoid organs following adoptive transfer in TCRα^-/-^ mice

To study the sufficiency of MAIT cells to provide B cell help in the context of mucosal immune responses *in vivo*, we utilized an adoptive MAIT transfer model into T*cra^tm1Mom^*/J (TCR*α*^-/-^) mice followed by a prime-boost model of mucosal bacterial infection. Expansion of lung MAIT cells from WT C57Bl/6 (B6) mice using an intranasal *Salmonella* Typhimurium challenge have shown to result in MAIT cells that retain function even after adoptive transfer (*40*). TCR*α*^-/-^ mice have been successfully used in T cell transfer experiments to study the B cell help potential of non-Tfh CD4 T cells without the potential masking effects of classic Tfh cells (*60*). We thus adoptively transferred expanded lung MAIT cells or PBS as an injection control (Sham groups) from B6 into TCR*α*^-/-^ mice and then utilized an intranasal prime-boost challenge model with live *Vibrio cholerae* O1 Inaba (*V.c*) that induces systemic and mucosal LPS IgA and IgG responses (*61*) (Fig. 4A). MAIT cells have been shown to be activated during cholera infection *in vivo* (*22*), and *V. cholerae* can biosynthesize riboflavin and thus can activate MAIT cells via MR1 presentation of riboflavin intermediates (*62*). CXCR5 expression was present but low on D7 *S.* Typhimurium expanded MAIT cells prior to adoptive transfer (fig. S10A-B). To confirm MAIT cells were successfully transferred into recipient mice, we measured MAIT frequencies and total cell numbers by flow cytometry of digested lung tissue and spleen (Fig. 4B and fig. S9A-C). The frequency and total cell count of lung MAIT cells at Day 42 in MAIT-*V.c* and MAIT-PBS groups were statistically similar to the WT-*V.c* group (Fig 4C and fig. S9B). Importantly, no MAIT cells were recorded in either Sham-*V.c* or Sham-PBS group. Similar MAIT cell counts were also observed in the spleen (fig. S9C), thus demonstrating WT physiologic MAIT frequencies and repopulation of various organs following adoptive transfer of lung MAIT cells. Furthermore, we found no differences between all TCR*α*^-/-^ groups in total non-MAIT TCR*β*^+^ T cells, thus confirming no significant expansion of contaminating non-MAIT T cells following adoptive transfer (fig. S9D). We next analyzed the expression of CXCR5 in CD8 T cells, CD4 T cells and MAIT cells from the WT-*V.c* and MAIT cells from the MAIT-*V.c* and MAIT-PBS groups (Fig. 4D and E). MAIT cells from WT-*V.c*, MAIT-*V.c* and MAIT-PBS groups, though they had decreased CXCR5 MFI compared to WT CD4 T cell populations, had significantly increased CXCR5 expression compared to the control CD8 T cell group in lungs, suggesting a potential mouse MAITfh population (Fig. 4D and E). Taken together, we found that adoptive transfer of expanded MAIT cells into TCR*α*^-/-^ mice repopulates MAIT cells, including CXCR5^+^ MAIT cells in lymphoid and mucosal organs, to levels similar to that of WT mice.

**Fig. 4.**
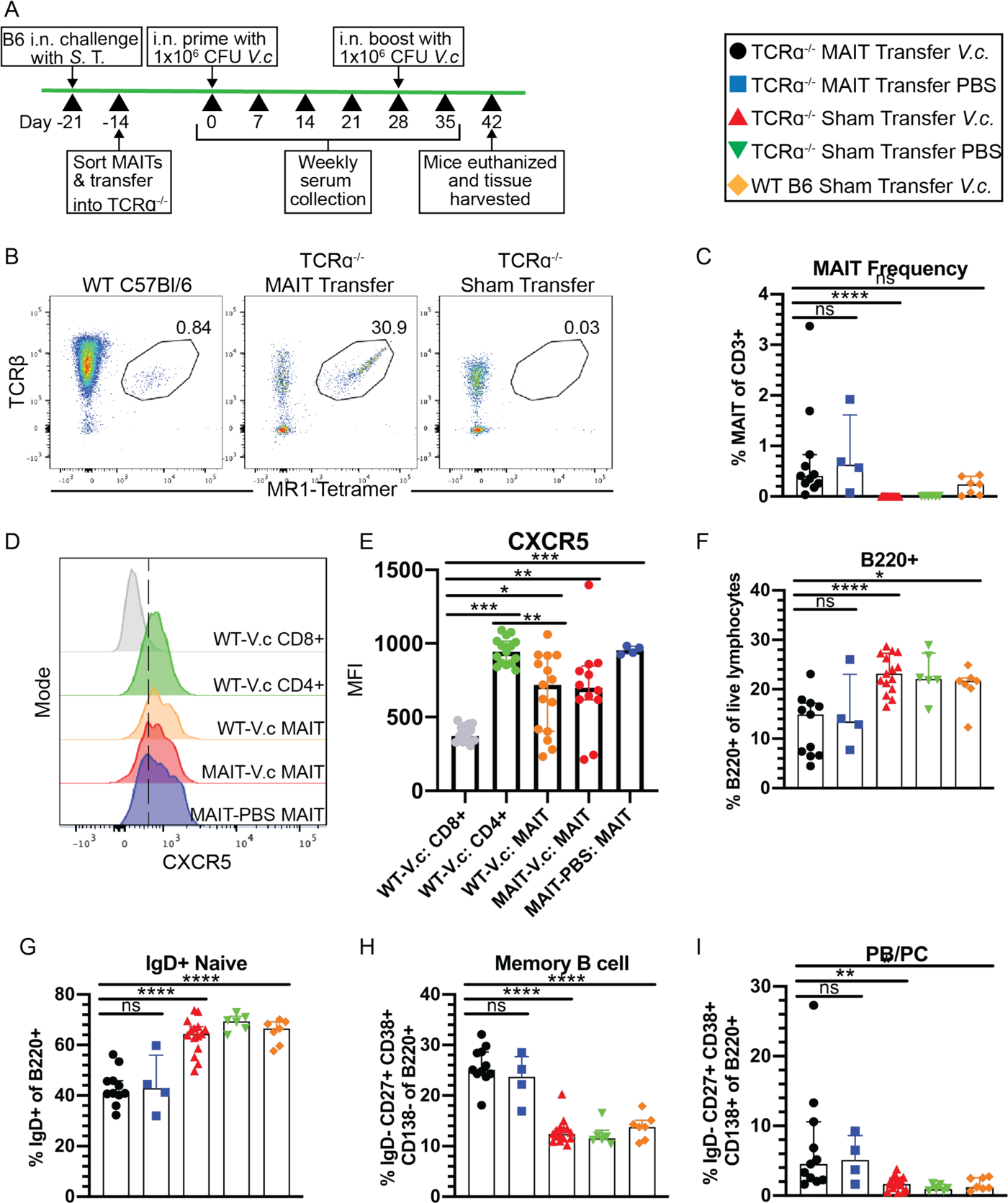
Adoptively transferred MAIT cells expand in lungs of TCR*α*^-/-^ mice driving B cell differentiation. (**A**) MAIT adoptive transfer and *V.c* intranasal challenge time-line. (**B**) Representative FACS plots of MAIT cells (TCR*β*^+^ MR1-Tetramer^+^) from lung tissue of WT-*V.c* (left), MAIT-*V.c* (middle), and Sham-*V.c* (right) mice at Day 42 post initial *V. cholerae* challenge. (**C**) MAIT cell frequency as percentage of CD3. Representative flow cytometry histograms (**D**) and Mean fluorescence intensity quantification (**E**) of CXCR5 in WT-*V.c* CD8 T cells (grey), CD4 T cells (green) and MAIT cells (orange), MAIT-*V.c* MAIT cells (red), and MAIT-PBS MAIT cells (blue). (**F**) Frequency of B220^+^ B cells among live lymphocytes. Frequency of IgD^+^ naïve B cells (**G**), IgD^-^ CD27^+^ CD38^+^ CD138^-^ Memory B cells (**H**) and IgD^-^ CD27^+^ CD38^+^ CD138^++^ PB/PCs (**I**) as percent of B220^low-high^ lymphocytes. Data are represented as Median with IQR from 4 independent experiments. n= 4-15 mice per group. *\*p* < 0.05, *\*\*p*<0.01, *\*\*\*p*<0.001, *\*\*\*\*p*<0.0001 by two-tailed Mann-Whitney *U* test. Legend in black box denotes experimental groups: MAIT-*V.c* = TCR*α*^-/-^ plus MAIT Transfer*-V.c* challenge, MAIT-PBS = TCR*α*^-/-^ plus MAIT Transfer*-*PBS challenge, Sham-*V.c* = TCR*α*^-/-^ plus Sham Transfer*-V.c* challenge, Sham-PBS = TCR*α*^-/-^ plus Sham Transfer*-*PBS challenge, WT-*V.c* = WT C57Bl/6J plus Sham Transfer-*V.c* challenge.

### MAIT cell adoptive transfer drives increase in plasmablast differentiation and memory B cell development

We next sought to determine the impact of MAIT adoptive transfer on mucosal B cell differentiation. Day 42 digested lung suspensions were analyzed by flow cytometry with B cells being defined as B220^+^ lymphocytes (fig S11A). Interestingly, total B220^+^ cell frequency (Fig. 4F) and total B cell count per lung (fig. S11B) were decreased in both the MAIT-*V.c* and MAIT-PBS groups compared to Sham-*V.c*, Sham-PBS and WT-*V.c* groups. Similar decreases in IgD^+^ naïve B cell frequency were observed in MAIT transfer groups (Fig. 4G), although no statistical differences were observed in total naïve B cells (fig. S11C). In contrast, we found that both MAIT-*V.c* and MAIT-PBS groups had higher Memory B cell (B220^+^ IgD^-^ CD27^mid^ CD38^+^ CD138^-^) and Plasmablast (PB)/Plasma cell (PC) (B220^low^ IgD^-^ CD27^+^ CD38^+^ CD138^++^) frequency as a percentage of total B cells compared to Sham-*V.c,* Sham-PBS and WT-*V.c* groups (Fig. 4H and I). Total Memory B cell count per lung was not statistically different between groups with significant variability observed in the MAIT-*V.c* group (fig. S11D), though we saw a trending increase in total PB/PC cell count per lung in MAIT-*V.c* compared to Sham-*V.c* (p=0.097) and WT-*V.c* (p=0.179) groups (fig. S11E). Overall, these data suggest that in the absence of *αβ* T cells, transfer of activated MAIT cells help induce B cell differentiation from naïve B cells to PB/PCs or memory B cells.

### Increase in V. cholerae specific IgA responses following MAIT transfer

*V.c*-specific antibodies in blood is a surrogate for protection against cholera (*63, 64*). Specifically, the vibriocidal assay, a measurement of complement-fixing bactericidal antibody activity, has been in use since the mid twentieth century (*65*) and shown to be the best marker of protection (*66, 67*). In addition to the vibriocidal titer, cholera toxin (CT) and lipopolysaccharide (LPS) serum ELISAs have been widely used to estimate cholera incidence (*67–70*). The O-specific polysaccharide (OSP) component of LPS and CT are the immunodominant antigens following cholera infection (*71*). Therefore, in order to measure the impact of MAIT transfer on pathogen-specific antibody kinetics we measured serum IgG, IgA, and IgM antibody responses against whole *V.c*-lysate, OSP, and CT. We found comparable *V.c*-lysate IgA responses in the MAIT-*V.c* and WT-*V.c* groups particularly following *V.c* re-challenge (D28), suggesting development of a memory response (Fig. 5A). Day 42 endpoint ELISAs reveal a significantly higher *V.c*-lysate IgA responses in the MAIT-*V.c* group compared to Sham-*V.c* (p<.0001), and MAIT-PBS (p<0.05) groups, and similar responses compared to the WT-*V.c* group (p=0.58) (Fig. 5C). Importantly, minimal *V.c*-lysate IgA responses were observed in the Sham-PBS group. MAIT-*V.c* and MAIT-PBS groups had no significant differences in *V.c-*lysate IgG (p=0.66, Fig. 5B and D) and IgM (p=0.08, fig. S12A and B) responses, despite higher compared to Sham transfer groups (p<0.01), thus indicating a non-specific response induced by MAIT transfer. We saw a non-significant (p=0.054) higher OSP IgA response in the MAIT-*V.c.* group compared to the Sham-*V.c* group, which can largely be attributed to high values in one mouse (Fig. 5E and G). No differences were observed in OSP-IgG responses between MAIT-*V.c* and Sham-*V.c* groups (p=0.64), Fig. 5F and H). Notably, even Sham-*V.c* TCR*α*-/- mice were able to mount (albeit lower than WT) OSP IgG responses, indicating that a lack of T cells blunted, but did not eliminate, the ability to mount an IgG response to a polysaccharide antigen, which is classically T-independent. In contrast, OSP-IgM levels were similar between the MAIT-*V.c* and Sham-*V.c* (p=0.33), and MAIT-*V.c* and WT-*V.c* groups (p=0.11), though were lower in the Sham-*V.c* compared to the WT-*V.c* (p=0.03) groups. All *V.c* infected groups had higher OSP-IgM responses compared to MAIT-PBS (p<0.05) and Sham-PBS (p<0.01) groups, indicating a specific but primarily T-independent response (fig. S12E and F). Overall, very low levels of CT IgA and IgG were observed in all TCR*α*^-/-^ groups (Fig. 5I-L) compared to the WT-*V.c* group, supporting the role of classical helper T cells in protein antigen B cell help, a response that adoptive transfer of MAIT cells did not rescue. Analysis of total serum IgA revealed a highly variable but substantial increase in the MAIT-*V.c* group over both Sham-*V.c* (p<0.0001)) and WT-*V.c* groups (p=0.01), fig. S12H). No statistically significant changes were observed in total IgM and IgG between MAIT-*V.c* and Sham-*V.c* groups (fig. S12G and I). To confirm the phenotype observed in the serum corresponded to the mucosa, Bronchoalveolar lavage (BAL) fluid from a subset of MAIT-*V.c* and Sham-*V.c* mice were assayed for *V.c* lysate IgA, IgG, and IgM responses. We found increased anti-lysate responses in all isotypes in the MAIT-*V.c* compared to the Sham-*V.c* group which corresponded with findings from serum responses (fig. S12J-L).

**Fig. 5.**
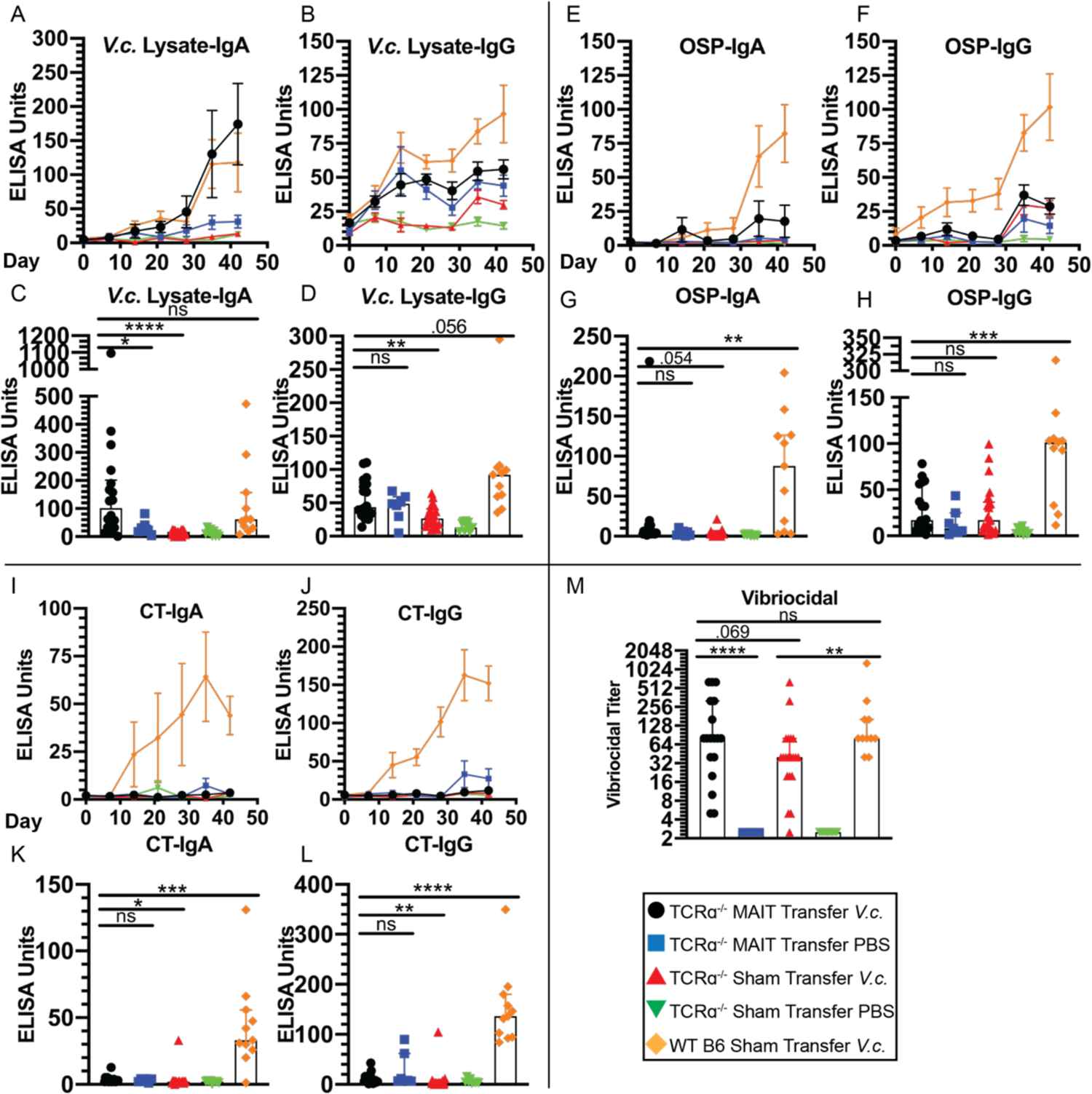
MAIT cells promote *V. cholerae*-specific IgA antibody responses in TCR*α*^-/-^ mice. Serum ELISAs against *V. cholerae* whole lysate, OSP and CT. (**A-D**) IgA and IgG *V. cholerae* lysate ELISAs from weekly cheek bleed (A & B) and Day 42 endpoint serum (C & D). (**E-H**) IgA and IgG OSP ELISAs from weekly cheek bleed (E & F) and Day 42 endpoint serum (G & H). (**I-L**) IgA and IgG CT ELISAs from weekly cheek bleed (I & J) and Day 42 endpoint serum (K & L). Data are represented as ELISA units normalized to a pooled serum positive control from WT B6 mice challenged with *V. cholerae*. (**M**) Vibriocidal titer of Day 42 endpoint serum. Data are represented as Median with IQR from 5 independent experiments. n= 6-22 mice per group. *\*p* < 0.05, *\*\*p*<0.01, *\*\*\*p*<0.001, *\*\*\*\*p*<0.0001 by two-tailed Mann-Whitney *U* test. Legend in black box denotes experimental groups.

We further utilized the vibriocidal assay to measure complement-mediated bactericidal activity, the best studied and most commonly used correlate of protection from cholera in humans and a proxy for protection in murine cholera challenge models (*66, 67, 72*). We found a trending increase in vibriocidal titer in the MAIT-*V.c* group compared to the Sham-*V.c* group (p=0.069), and no difference when compared to the WT-*V.c* group (Fig. 6M). Unlike the ELISAs, no background vibriocidal responses were observed in the MAIT-PBS group, demonstrating the specificity of this assay. Taken together, adoptive transfer of MAIT cells into T-cell deficient mice rescued deficiencies in *V. cholerae*-specific IgA and promoted slight increases in vibriocidal antibody responses.

## Discussion

MAIT cells are widely appreciated for their ability to respond rapidly to microbial antigens and cytokines through potent production of Th1 and Th17-associated inflammatory cytokines and cytotoxic molecules. Recent studies have highlighted the capacity of human MAIT cells to provide B cell help *in vitro*, and promote B cell differentiation and antibody production in animal models (*23, 25, 26*). Despite these findings, whether there remains a pre-defined MAIT subset capable of B cell help remains uncertain.

Additionally, many studies have detected MAIT cells within mucosal lymphoid tissues but have yet to understand their role in these environments (*33, 41, 73*). In this study we begin to define the role of MAIT cells in human mucosal lymphoid organs and uncover a novel T follicular helper-like MAIT population defined by CXCR5. This MAITfh subset highly expresses PD1, ICOS, *BCL6*, and IL-21, and preferentially locates near germinal centers. We further add to the body of literature proving the ability of MAIT cells to provide B cell help *in vivo* through sufficiency experiments following mucosal challenge. Here we show that adoptively transferred MAIT cells can rescue microbe specific IgA responses, and promote B cell differentiation in the absence of other *αβ*T lymphocytes.

We report a subset of CXCR5^+^ MAIT (MAITfh) cells that are enriched in mucosal lymphoid organs and represent a small percentage of peripheral blood MAIT cells (Fig. 1C). We further report that MAITfh cells highly express the B cell help co-stimulatory molecules, PD1 and ICOS, compared to CXCR5^-^ MAIT cells (Fig 1D and E). In addition, we show high relative expression (*≍*30-45 fold) of *CD40L* in both MAIT and MAITfh groups compared to the control CD8+ population (fig. S5B). PD1, ICOS and CD40L expression are a hallmark of GC Tfh cells (*74*). PD1, although long considered a marker of T cell exhaustion, has an important role in Tfh GC positioning and function (*5*). ICOS and CD40L engagement with ICOS Ligand and CD40 expressed on B cells is essential for GC formation and Tfh activation (*6, 7*). CD40L is also implicated in MAIT induced maturation of dendritic cells (*75*). Based on the Tfh costimulatory molecule profile, we hypothesized that MAITfh cells may share further transcriptional similarities with Tfh cells. In support, we found that MAITfh cells in PBMCs have higher *BCL6* expression than MAIT and Tfh cells and a trending increase in *BCL6* expression in tonsils (Fig. 2E).

BCL6 is considered the master regulator of the Tfh lineage (*76*) and is similarly required for iNKTfh B cell help responses (*18*). Although further research into the transcriptional regulation of MAITfh cells is necessary to understand the requirements for differentiation, plasticity and function, these data support a B cell help transcriptional phenotype among the MAITfh subset.

We report MAITfh expression of the B cell help cytokines *IL-21*, *IL-10*, *IL-6* and *BAFF* (Fig. 2 and fig. S3). Specifically, we found trending increases in *IL-21* and *IL-6* expression in MAITfh compared to MAIT cells in tonsils, while *BAFF* and *IL-10* were highly expressed in both PBMC and tonsil MAIT cells compared to Tfh controls (Fig. 2). IL-21 plays an important role in T dependent B cell differentiation, class switching and antibody production (*77*) and together with IL-6 promotes Tfh differentiation (*78*).

IL-10 promotes antibody production and class switching (*79*). IL-21 expression in MAIT cells has been demonstrated in low concentrations in PBMCs (*23*), and in a subset of PD1^High^ CXCR5^-^ MAIT cells in pleural effusions from Tuberculosis patients (*24*). This study also presents a small subset of CXCR5^+^ PD1^high^ MAIT cells, though they do not investigate whether this population expressed IL-21 as well (*24*). In contrast, we show that tonsil CXCR5^+^ MAIT cells have a higher frequency of IL-21^+^ IFN*γ*^-^ cells (Fig. 2J) and produce more IL-21 than CXCR5^-^ MAIT cells (Fig. 2K). It is unclear whether PD1^High^ CXCR5^-^ MAIT cells in pleural effusions are transcriptionally or developmentally related as potential precursors to CXCR5^+^ MAIT cells found in lymphoid tissues. Furthermore, the B cell survival cytokine, BAFF, is integral in T independent class switching and differentiation (*80*), and is similarly produced by iNKTfh cells to promote B cell responses (*18*). Thus, it is possible MAITfh cells can aid in B cell help through cognate interactions via direct MR1-TCR, co-stimulatory molecule engagement and cytokine expression, or through non-cognate interactions such as promoting Tfh differentiation or licensing of dendritic cells.

We have observed that PD1^high^ MAIT cells preferentially locate within or around GCs in human tonsils (Fig. 3A and B). This is noteworthy since spatial positioning of T and B cells within lymphoid tissues is critical in understanding their phenotype. For instance, CXCR5^+^ PD1^High^ GC Tfh cells reside primarily in the light zone of GCs where they promote affinity maturation and differentiation of GC B cells leading to high affinity antibody responses (*74*). Thus, by combining imaging results with *in vitro* MAITfh expression of B cell help co-stimulatory markers and cytokines (Fig. 1 and 2), we can conclude that MAITfh cells share both phenotype and positioning with traditional GC Tfh cells. Thus we hypothesize that this population may be involved in cognate or contact-dependent B cell help as is seen in other innate-like Tfh subsets, such as iNKTfh (*12*). Though as this population is relatively small in tonsils, further research will need to address the *in vivo* relevance to overall mucosal antibody production, the role of MAITfh-B cell interactions in affinity maturation, and development of B cell memory.

We report that adoptive transfer of murine MAIT cells boosts pathogen-specific antibody responses in the absence of other *αβ*T lymphocytes. We further previous work on MAIT-B cell help responses in animal models (*25, 26*), by using an adoptive transfer model to show that MAIT cells are sufficient to increase functional anti-bacterial antibodies in the context of a mucosal challenge. Most notably, we show that adoptive transfer of MAIT cells rescues the deficiency in *V.c-*specific IgA of infected TCR*α*^-/-^ mice when compared to infected WT mice. Examination of antibody kinetics of *V.c* specific IgA in mice receiving MAIT transfer suggest a memory phenotype, based on substantial increases on Day 35 (one-week post re-challenge) (Fig. 5A), and is associated with higher lung PB/PC and memory B cell frequencies (Fig. 4H and I). Interestingly, we also report MAIT-*V.c* mice had significant systemic *V.c* specific IgA and total IgA responses compared to Sham-*V.c* mice (Fig. 5A and fig. S12H). These data may be a result of the mucosal nature of the intranasal challenge, which has shown to induce both mucosal and systemic immunity in other bacterial infections (*81*). Notably, increases in total IgA in MAIT-PBS mice suggests MAIT cells may specifically promote IgA class switching. In support, pulmonary MAIT cells in mice have recently been shown to produce both IL-10 (*82*) and TGF1-*β* (*83*), key cytokines in IgA class switching (*84*). Although CXCR5 staining of lung MAIT cells suggests a similar Tfh-like MAIT phenotype in mice (Fig. 4D and E), limitations of available reagents for identifying murine MAIT cells by IHC precludes our ability to confirm a MAITfh subset in mice. Future analysis of lung and lymph node MAIT cells is necessary to confirm a MAITfh subset in mice and its potential role in IgA class switching and antibody production. Given that systemic *V. cholerae* specific IgA and vibriocidal responses strongly correlate with protection against subsequent cholera re-infection (*64, 66, 67*), these are promising data in the context of mucosal vaccines, where IgA is often the primary means of protection, and development of long-lasting protective memory responses is limiting (*85*). These data support further research into the potential adjuvant activity of MAIT ligand administration during mucosal infection or vaccination to enhance humoral immune responses.

In addition to promoting microbe-specific IgA responses, we report amplification of non-specific antibodies following MAIT transfer. We report similar *V. cholerae* lysate IgG (Fig. 5B and D) and IgM (fig. S11A and B), OSP (Fig. 5E-H), CT (Fig. 5I-L), total IgA (fig. S11H), and similar changes in B cell subset frequencies in our MAIT-PBS and MAIT-*V.c* groups (Fig. 4F-I). Thus, regardless of *V. cholerae* infection, adoptively transferred MAIT cells are promoting B cell differentiation and antibody production. Similar MAIT amplification of non-specific responses were reported in non-human primates (*26*) and Fc*γ*RIIB^-/-^ mice (*25*). Specifically, Rahman et al. reported increased total IgM and IgG antibody production and increased IgD^-^ B cells (indicating class switching) when non-human primate MAIT cells were co-cultured with B cells. Murayama et. al 2019 show increases in total IgG and anti-dsDNA IgG and IgA when MAIT cells were co-cultured with B cells from Fc*γ*RIIB^-/-^ mice. They also report reduction in anti-dsDNA IgG in MR1^-/-^ (MAIT deficient) Fc*γ*RIIB^-/-^ mice, though total IgG was not published. It should be noted that in all studies, MAIT cells were either activated *in vitro* in direct co-culture, or *in vivo* in the absence of regulatory T cells (TCR*α*^-/-^ mice). Therefore, such responses may be a result of lack of either MAIT and/or B cell regulation leading to aberrant antibody responses. This has interesting consequences in the setting of mucosal immunity and in autoimmunity. In a setting of proper T and B cell regulation, MAIT cells may help to promote anti-microbial responses and therefore could potentially be targeted as a vaccine adjuvant. Conversely, in a dysregulated environment, such as in Fc*γ*RIIB^-/-^ or TCR*α*^-/-^ mice, MAIT cells may enhance aberrant antibody production against self or commensal microbes thus enhancing the pro-inflammatory autoimmune state. These aberrant responses open many questions into cell extrinsic MAIT regulation which remains relatively unstudied.

There are a number of limitations that need to be addressed when considering the results of this study. In our human *in vitro* experiments, we primarily compare PBMCs from healthy adults ages 18+ to tonsils from pediatric patients, ages 2-16, undergoing tonsillectomy. MAIT cells expand immediately after birth and reach adult frequencies by approximately 10 years of age, and most reach maturity (CD45RO^+^) by 6 months of age (*86*). Therefore, it is possible that age may explain low MAIT percentage in tonsil samples, though analysis of 5 paired PBMC and tonsil samples pediatric donors revealed little differences in MAIT frequency when compared to the healthy donor adult PBMCs (fig. S4). Limited MAIT frequency in tonsils also limited our abilities to perform mechanistic analysis of our MAITfh populations. Future co-culture studies with B cells and expanded MAITfh cells may be useful to understand the MAIT-B cell help mechanism. Regarding our animal studies, a limitation of this study is the under sampled analysis of mucosal antibody measurement, as we focused primarily on systemic responses in serum. While mucosal secretory IgA (sIgA) and circulating IgA responses may differ, measurement of systemic antibodies is standard practice for mucosal vaccine response testing in humans, and *V.c* specific serum IgA response and vibriocidal titer strongly correlate with protection in humans (*64, 66, 67*). Additionally, our limited analysis of BAL antibody responses correlated well with serum results. Thus, these serum responses are likely relevant to mucosal protection, though lack of conclusive protective immunity via survival curves or colony counts limits this finding. Secondly, our study lacks insight into the mechanism and location of MAIT B cell help in our murine model. Future studies, potentially using congenic markers, are necessary to address whether MAIT cells provide help within local lymph nodes, MALT or within the mucosal lamina propria. Additionally, the nature of *the V. cholerae* antigens targeted by the IgA antibodies remains unknown. Despite previous evidence showing association of human MAIT cells with polysaccharide specific antibody responses (*22, 35*), we found only moderate changes in OSP antibody responses in our MAIT transferred mice, suggesting an alternate antigen target. Future studies are necessary to address the whether MAIT cells, like other innate-like T cells, promote antibody responses to specific antigen targets.

In conclusion, we identify a Tfh like MAIT population enriched near germinal centers of mucosal lymphoid tissues that express cytokine and transcriptional profiles consistent with B cell help. In addition, we show in an animal model that MAIT cells are sufficient to rescue pathogen-specific IgA antibody responses following mucosal challenge in the absence of other classical T cells. Our work strengthens a growing body of evidence supporting the capacity of MAIT cells to aid in B cell help and antibody mediated immunity, and support the expanding phenotypic heterogeneity of MAIT cells (*87*). As MAIT cells are enriched at the mucosa and can be activated with specific ligands, we hypothesize that they may be a promising target to enhance mucosal vaccines.

## Materials and Methods

### Peripheral blood and tonsil collection

Healthy adult blood was collected from residual leukocyte packs following blood donation (ARUP, Sandy, UT), and PBMC’s were isolated by density gradient centrifugation (Lymphoprep, StemCell Technologies). Residual palatine tonsil samples were acquired from pediatric patients, ages 2-16, undergoing routine tonsillectomies for tonsillar hyperplasia or recurrent tonsillitis. All tonsil samples were stored on ice in complete media (RPMI 1640 with 10% FBS, 1% penicillin/streptomycin, and 15 mM HEPES) (R10) and processed within 2 hours of surgery. 1 cm x 1 cm pieces were cut and frozen in O.C.T. Compound (Fisher) at −80°C for later use. The remaining tonsil tissue was minced in media and dissociated using a 100 μM cell strainer. Tonsil mononuclear cells were isolated by density gradient centrifugation (Lymphoprep, StemCell Technologies). PBMC’s from pediatric donors were obtained during surgery with informed consent granted by a parent/guardian. PBMC and tonsil mononuclear cells were frozen in media containing 25% FBS and 10% DMSO at −80°C for later use. All human samples were de-identified prior to receipt, and the research protocol was deemed exempt by the University of Utah Institutional Review Board (Protocol 100683).

### Flow cytometry and cell sorting

Prior to surface staining, human and mouse single cell suspensions were labeled with Fixable Viability Dye eFluor 780 (eBioscience) according to manufacturer’s protocol to delineate live cells. Cells were washed and incubated with fluorochrome conjugated antibodies for 20 min at room temperature (RT). Mouse cells were incubated in anti-mouse CD16/CD132 (Fc block) for 15 minutes prior to surface staining. Mouse and human MR1-Tetramers (NIH Tetramer Core) diluted 1:400 were added to surface stain. Prior to mouse MR1-Tetramer staining, cells were incubated for 15 minutes with unlabeled i6FP-Tetramer (NIH Tetramer Core) to minimize non-specific binding. For intracellular cytokine staining, cells were fixed and permeabilized using Foxp3/Transcription Factor Staining Buffer set (eBioscience) according to manufacturer’s protocol and incubated with fluorochrome conjugated antibodies for 40 min at RT. Cells were analyzed using BD LSR Fortessa or Cytek Aurora for phenotypic analyses, or Aria II (BD Biosciences) for cell sorting. The following monoclonal antibodies to human were used: CD3-AF700 (clone OKT3), CD8-FITC/BV605 (cloneSK1, TCR V*α*7.2-BV711/PE-Cy7 (clone 3C10), PD1-BV605/AF700 (clone EH12-2H7), ICOS-PerCP-Cy5.5/BV605 (clone C398.4A), CXCR5-PE-Cy7/APC (clone J252D4), IL-21-APC (clone 3A3-N2), IFN*γ*-BV421 (clone B27), IL-6-FitC (clone MQ2-13A5), IL-10-PerCP-Cy5.5 (clone JES2-9D7), CD161-AF488 (clone HP-3610) (Biolegend), CD4-BUV496 (clone SK3), CD3-BUV395 (clone SK7) (BD Biosciences). The following monoclonal antibodies to mouse were used: CD3-PE-Dazzle594 (clone 17A2), TCR*β*-BV421 (clone H57-597), CXCR5-BV605 (clone L138D7), CD44-BV650 (clone IM7), TCR*γδ*-PE-Cy7 (clone GL3), CD8-APC (clone 53-6.7), CD45R-PE-Cy5 (clone RA3-6B2), IgD-BV510 (clone 11-26c.2a), CD138-BV605 (clone 281-2), CD38-PE-Cy7 (clone 90, CD27-APC (clone LG-3A10) (Biolegend), CD4-FITC (clone GK1.5) (Tonbo Biosciences).

### In vitro stimulation assays

For flow cytometry intracellular cytokine staining, 1-2 x 10^6^ mononuclear cells isolated from blood or tonsil samples were cultured in a round bottom 96 well plate at 37°C 5% CO2 in R10 media for 6 hours with or without 200 ng/ml phorbol myristate acetate (PMA) and 1.0 μg/ml Ionomycin (Sigma-Aldrich) or 0.5 μM/ml 5-amino-6-D-ribitylaminouracil (5-A-RU) (courtesy of Jeffrey Aubé, UNC) and 50 μM/ml Methylglyoxal (Sigma). Brefeldin A Solution (Biolegend) was added at 1X for the final 4 hours. For measurement of IL-21 production, tonsil mononuclear cells were sorted into CD8^+^ CXCR5^-^, CD4^+^ CXCR5^+^ PD1^High^, MR1-Tet^+^, V*α*7.2^+^, CXCR5^-^, MR1-Tet^+^, V*α*7.2^+^, CXCR5^+^ populations and 4×10^4^ cells in R10 media were cultured in a round bottom 96 well plate for 48 h with 20 ng/ml PMA and 1.0 μg/ml Ionomycin as described in Shen et al (*55*). Plates were centrifuged and cell supernatant was isolated for ELISAs.

### qRT-PCR

PBMC and tonsil CD8^+^ CXCR5^-^, CD4^+^ CXCR5^+^, MR1-Tet^+^ V*α*7.2^+^ CXCR5^-^, MR1-Tet^+^ V*α*7.2^+^ CXCR5^+^ cells were sorted into TRIzol Reagent (Qiagen). Total RNA was isolated using the RNeasy Micro kit and cDNA synthesized a High-Capacity cDNA Reverse Transcription Kit (Applied Biosystems). Targeted qPCR was performed using the QuantStudio 6 Real-Time PCR system using the following Taqman Primers (Applied Biosystems): BCL6 (Hs00153368_m1), Tnfsf13b (Hs00198106_m1), IL6 (Hs00174131_m1), IL10 (Hs00961622_m1), IL21 (Hs00222327_m1), TNF*α* (Hs00174128_m1), TGFB1 (HS00998133_m1), Tcf7 (Hs01556515_m1), Tbx21 (Hs00894392_m1), Runx3 (Hs01091094_m1), Rorc (Hs01076112_m1), Perforin (Hs00169473_m1), Pdcd1 (Hs01550088_m1), Il17a (Hs00174383_m1), Ifn*γ* (Hs00989291_m1), Icos (Hs00359999_m1), Gzmb (Hs00188051_m1, Cxcr5 (Hs00173527_m1), Cd40l (Hs00163934_m1), Cd38 (Hs01120071_m1), Blimp1 (Hs00153357_m1), Ascl2 (Hs00270888_s1), ActB (Hs99999903_m1).

### Single cell TCR sequencing

MAITs from PBMC’s and tonsils were single cell sorted using the Aria II cell sorter (BD Biosciences) directly into One Step RT-PCR reaction mix (NEB) loaded in MicroAmp Optical 96-well reaction plates (Applied Biosystems). MAITs were defined as CD3^+^ V*α*7.2^+^ MR1-Tetramer^+^ cells and sorted based on CD161 expression. Following reverse transcription and preamplification reaction, a series of three nested PCR’s were run using primers for TCR sequence and gene expression as described (*45*). Subsequent sequencing data analysis was performed as described (*45*).

### ELISAs and Vibriocidal assay

IL-21 concentration from tonsil mono-cultures was measured using Human IL-21 ELISA MAX Deluxe kit (Biolegend) according to manufacturer’s protocol. Serum *V.c* Lysate, OSP:BSA and CT ELISAs were performed as follows. Nunc Maxisorp flat bottom 96 well plates (Invitrogen) were coated with *V.c* Lysate or OSP:BSA (gift from Dr. Edward Ryan (Massachusetts General Hospital, Boston, MA) (1.0 μg/ml) in PBS and incubated overnight at 4°C. For CT ELISAs, Nunc plates were first coated with Monosialoganglioside GM1 (Sigma-Aldrich) (1.0 μg/ml) in 50mM carbonate buffer overnight at 4°C. Plates were subsequently blocked with 1% bovine serum albumin (BSA) (Sigma-Aldrich) and incubated overnight with CT (Sigma-Aldrich) (2.5 μg/ml) in carbonate buffer. All plates were then blocked with 1% BSA and 25 μl of mouse serum diluted 1:10 in 0.1% BSA-0.5% Tween 20 in PBS (BSA-PBST) was added for 2 hours. Plates were incubated with anti-mouse IgG (Invitrogen), IgA (Invitrogen), or IgM (Life Technologies) HRP conjugate diluted 1:1000 in BSA-PBST for 2 hours. 100 μl of TMB substrate (Thermo Fisher) was added and plates were immediately read kinetically at 405 nm (7 min x 1 min intervals) using plate reader (Biotek). ELISA measurements were recorded as max interval slope and normalized to pooled positive mouse sera included on each plate. Total Ig ELISAs were performed as described above except Nunc plates were pre-coated with anti-mouse IgA (clone RMA1), anti-mouse IgG (clone Poly 4053), or anti-mouse IgM (clone RMM-1) (Biolegend) and standard curves were generated using 2-fold dilutions of purified Mouse IgA, IgM and IgG (Southern Biotech). Plates were incubated with corresponding anti-mouse HRP antibodies and TMB substrate was neutralized after 10 min with 0.2 N HCl. Plates were read at 605 nm and concentrations were calculated based on the standard curve.

Vibriocidal titer was measured as described (*88*). *Vibrio cholerae* serotype 01 Strain El Tor Inaba N16961 grown in LB media was used as target organism for assay, and serum samples were diluted to a starting concentration of 1:5.

### Immunofluorescence staining

Unfixed pediatric tonsil samples frozen in OCT medium were cut into 9μm thick sections using a cryostat (Leica) and stored at −80°C. Tonsil sections were brought to room temperature and then fixed at −20°C for 10 min in pre-cooled 50:50 Acetone:Methanol solution. Sections were dried and then rehydrated in PBS for 5 min with gentle shaking. The sections were then blocked in 2% normal mouse serum (Invitrogen) for 1 hour and subsequently stained overnight at 4°C with mouse anti-human TCR V*α*7.2-PE (Clone 3C10, Biolegend), mouse anti-human CD161-AF488 (Clone NKR-P1A, Biolegend), mouse anti-human IgD-BV421 (Clone IA6-2, Biolegend) and mouse anti-human PD1-APC (Clone EH1202H7, Biolegend) all diluted 1:100. Sections were washed 3x in PBS and mounted using ProLong Gold Antifade Mounting media (Thermo Fisher). Full tonsil cross section images were acquired using the Leica Sp8 Confocal microscope and then processed and analyzed using ImageJ.

### Bacterial culture and V. cholerae lysate preparation

The attenuated *Salmonella* Typhimurium strain BMM50 was a gift from Dr. Stephen McSorely (UC Davis, Davis, CA). BMM50, like previously published *S.* Typhimurium BRD509 (*39*), has a deleted *aroA* gene and intact riboflavin pathway. *Vibrio cholerae* serotype 01 Strain El Tor Inaba N16961 was a gift from Dr. Edward Ryan (Massachusetts General Hospital, Boston, MA). All strains were inoculated from single colonies into Luria Bertani (LB) broth shaking at 220 RPM 37°C overnight. They were then subcultured 1:10 in fresh LB and incubated for 3-4 hours shaking at 37°C until log phase was reached.

Cultures were washed 2x in sterile PBS and normalized to 0.1 OD600 using a microplate absorbance spectrophotometer (BioRad). *V. cholerae* at 0.1 OD600 was used as inoculum for intranasal challenge model. *S.* Typhimurium BMM50 was diluted 1:10 with PBS to an OD600 of 0.01 for adoptive transfer inoculum. Inoculums ranged from 1×10^6^-1×10^7^ colony forming units (CFU) confirmed by plate counts. *V. cholerae* lysate for ELISAs was prepared as follows. *V. cholerae* was cultured from a single colony in LB broth shaking at 220 RPM 37°C overnight. *V. c* culture was then pelleted and washed 3x with cold sterile PBS. Following resuspension in PBS, sample was sonicated at 40% power 4 x 1 min intervals, keeping on ice for 1 minute between runs. Sonicated sample was then centrifuged at 15000 rpm x 10 min at 4°C, and the supernatant removed and sterilized using 0.2 μm syringe filter (Thermo Fisher). Protein content was measured using 260/280 absorbance ratio with Take3 Micro-Volume Plate reader (BioTek)

### Mice

T*cra^tm1Mom^*/J mice (*89*) (on a B6 background) were obtained from The Jackson Laboratory (Bar Harbor, ME) and bred at the University of Utah Comparative Medicine Center mouse facility. WT C57BL/6J mice were obtained directly from The Jackson Laboratory prior to the start of the experiment. 6-10-week-old female and male mice were used in adoptive transfer studies. All mice were housed under specific pathogen-free conditions. Experiments were performed in strict accordance with the NIH guide for the Care and Use of Laboratory animals and institutional guidelines for animal care at the University of Utah under approved protocol #17-01011.

### Adoptive transfer

Adoptive transfer of pulmonary MAIT cells was adapted from protocol described (*40*). C57BL/6 mice were infected i.n. with 10^6^ CFU *S.* Typhimurium BMM50 suspended in 50 μl sterile PBS (25 μl per nares). Mice were rested for 7 days to allow for MAIT expansion, in which mice were weighed daily and evaluated for signs of clinical pneumonia symptoms (respiratory distress, inactivity, ruffled fur, hunched posture). Mice with loss of >20% of body weight and/or severe clinical symptoms were euthanized. On day 7, mice were euthanized and the lungs were perfused with 10 mls of PBS through the heart and then removed. Single cells suspensions of lungs were prepared using the gentleMacs Lung Dissociation kit, mouse (Miltenyi Biotech) according to the manufacturer’s protocol. Red blood cells were lysed using ACK Lysis buffer (Thermo Fisher), and single cell suspensions were stained and sorted using the BD FACS Aria II. MAIT cells were defined as live CD3^+^ B220^-^ TCR*γδ*^-^ CD44^high^ TCR*β*^+^ MR1-Tetramer^+^ lymphocytes. Approximately 5×10^4^ - 1×10^5^ MAIT cells were sorted per mouse. 10^5^ MAIT cells suspended in 100 μl was transferred via retroorbital injections into lightly anesthetized mice. Transferred mice were monitored to confirm recovery following injections, and rested for 2 weeks to allow for MAIT expansion before use in intranasal challenge model.

### Intranasal challenge model

The following protocol was adapted from Nygren et al (*61*) to induce *V. cholerae* specific antibody responses through a prime-boost live bacterial model. In brief, on Day 0 mice were anesthetized using isoflurane and infected i.n. with 10^6^-10^7^ CFU *V. cholerae* 01 Biotype El Tor Serotype Inaba suspended in 50 μl sterile PBS (25 μl per nares). Mice were monitored for 7 days post infection for weight loss and clinical signs of pneumoniae (outlined above). Mice with loss of >20% of body weight and/or severe clinical symptoms were euthanized. Mice were subsequently re-challenged on Day 28 using the above protocol. Blood samples were collected weekly, prior to infections on Day 0 and 28, from submandibular bleeds on lightly isoflurane anesthetized mice. Blood serum was isolated using BD Microtainer SST tubes (BD Biosciences) according to the manufacturer’s protocol. On Day 42 (2 weeks post re-challenge), mice were euthanized using a bell jar and isoflurane. Blood was collected immediately via cardiac puncture and serum isolated. Lungs were perfused and single cell suspensions were processed as described above.

Spleens were excised, ground through 70 μm filters in cold R10 media. Red blood cells were lysed using ACK Lysis Buffer (Thermo Fisher). 1-2×10^6^ cells from lung and spleen cell suspensions were stained with fluorochrome conjugated antibodies for flow cytometric analyses.

### Statistical analysis

All statistical tests were performed using Prism version 8.4.2 (GraphPad Software, La Jolla, CA, USA). Differences were compared using two-tailed Mann Whitney *U* tests, or Multiple t tests comparisons accounting for False Discoveries using the Benjamini, Krieger and Yekutieli correction as indicated. Graphs were created using Prism or R (Vienna, Austria) and the graphing package ggplot2 (H. Wickham, Springer-Verlag, NY, USA).

## Author Contributions

O.J. and D.T.L. designed and directed the project. O.J and S.T. performed the experiments. K.F. and J.S.H. contributed to experimental methodology. J.D.M. provided samples and resources. D.T.L acquired funding. O.J. and D.T.L wrote the paper, and all authors discussed results, provided critical feedback, and contributed to the final manuscript.

## Supplementary Materials

**Supplementary Fig. 1.**
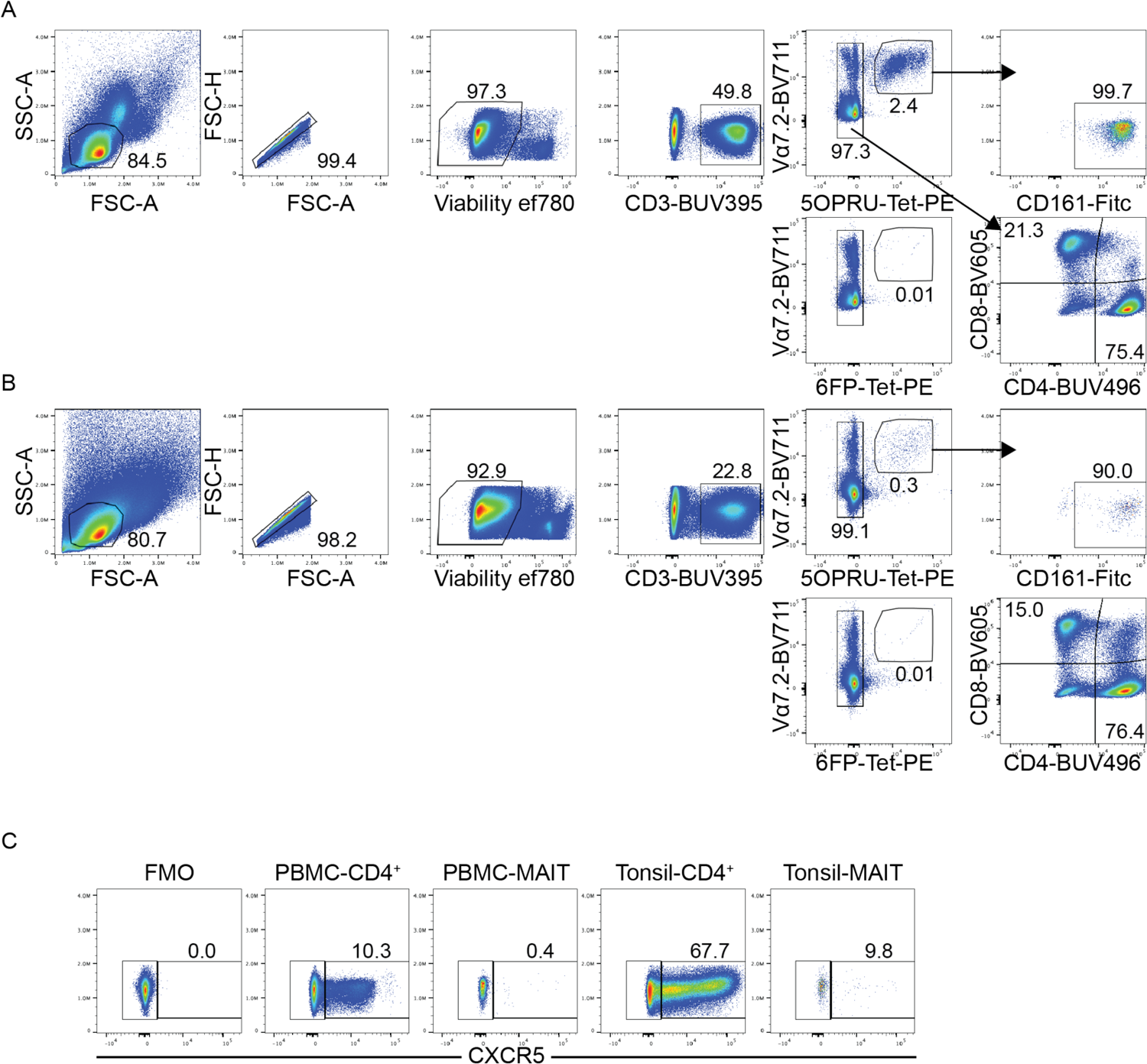
Representative flow cytometry gating strategy for Fig. 1. Representative flow cytometry gating strategy for MAIT and non-MAIT T cells in (**A**) PBMC and (**B**) tonsils. PE conjugated 6-FP MR1-Tetramer gating controls for PBMC and tonsils are located below 5-OP-RU MR1-Tetramer staining gates. (**C**) Representative CXCR5 staining for PBMC and tonsil CD4^+^ T and MAIT cells quantified in Fig. 1C. Gating was based on CXCR5 fluorescence minus one (FMO) control.

**Supplementary Fig. 2.**
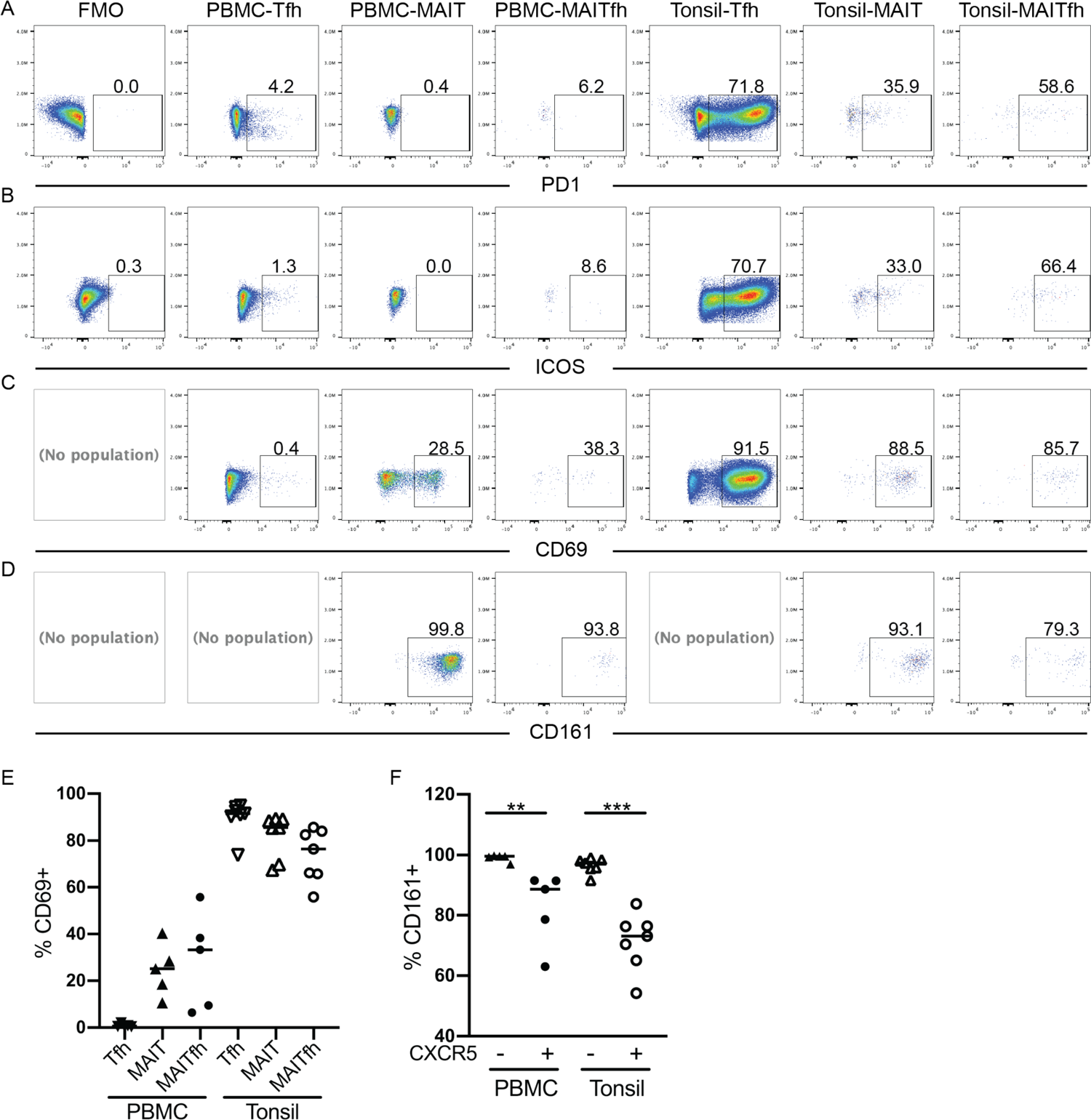
Flow gating strategy for T cell co-stimulatory molecule expression analysis. Representative flow cytometry gating strategy for PBMC and tonsil Tfh, MAIT and MAITfh cell (**A**) PD1, (**B**) ICOS, (**C**) CD69, and (**D**) CD161 expression. FMO’s were not included for CD69 and CD161 due to clarity of antibody staining. (**E**) Frequency of Tfh, MAIT, and MAITfh cell populations expressing CD69. (**F**) Frequency of CD161^+^ MAIT cells broken down by CXCR5 expression. Data are represented as median from 1 independent experiment. n = 5-7. *p < 0.05, **p<0.01, ***p<0.001, ****p<0.0001 by two-tailed Mann-Whitney U test.

**Supplementary Fig. 3.**
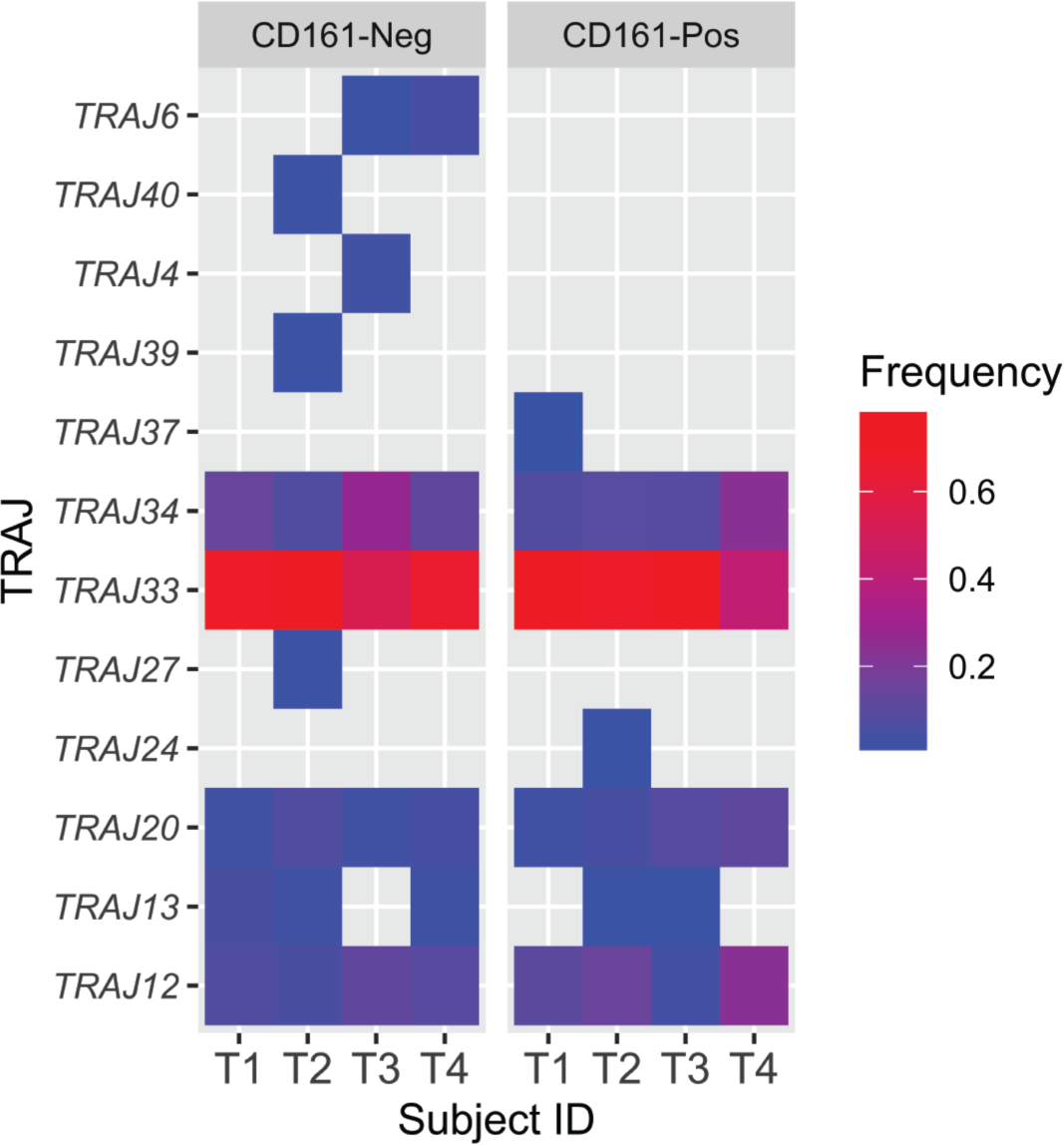
Similar TRAJ expression in CD161+ and CD161-tonsil MAIT cells. Heat map of TRAJ usage percent frequencies based on Illumina single cell TCR-Phenotype sequencing in tonsil MAIT cells broken down by CD161 negative (top) and positive (bottom) populations. Each column represents an individual donor (n=4). MAIT cells were defined as MR1-5-OP-RU-tetramer positive, and TRAV1-2 antibody positive. No statistical significance was found between CD161- and CD161+ groups using Multiple t tests comparison accounting for False Discoveries using the Benjamini, Krieger and Yekutieli method.

**Supplementary Fig. 4.**
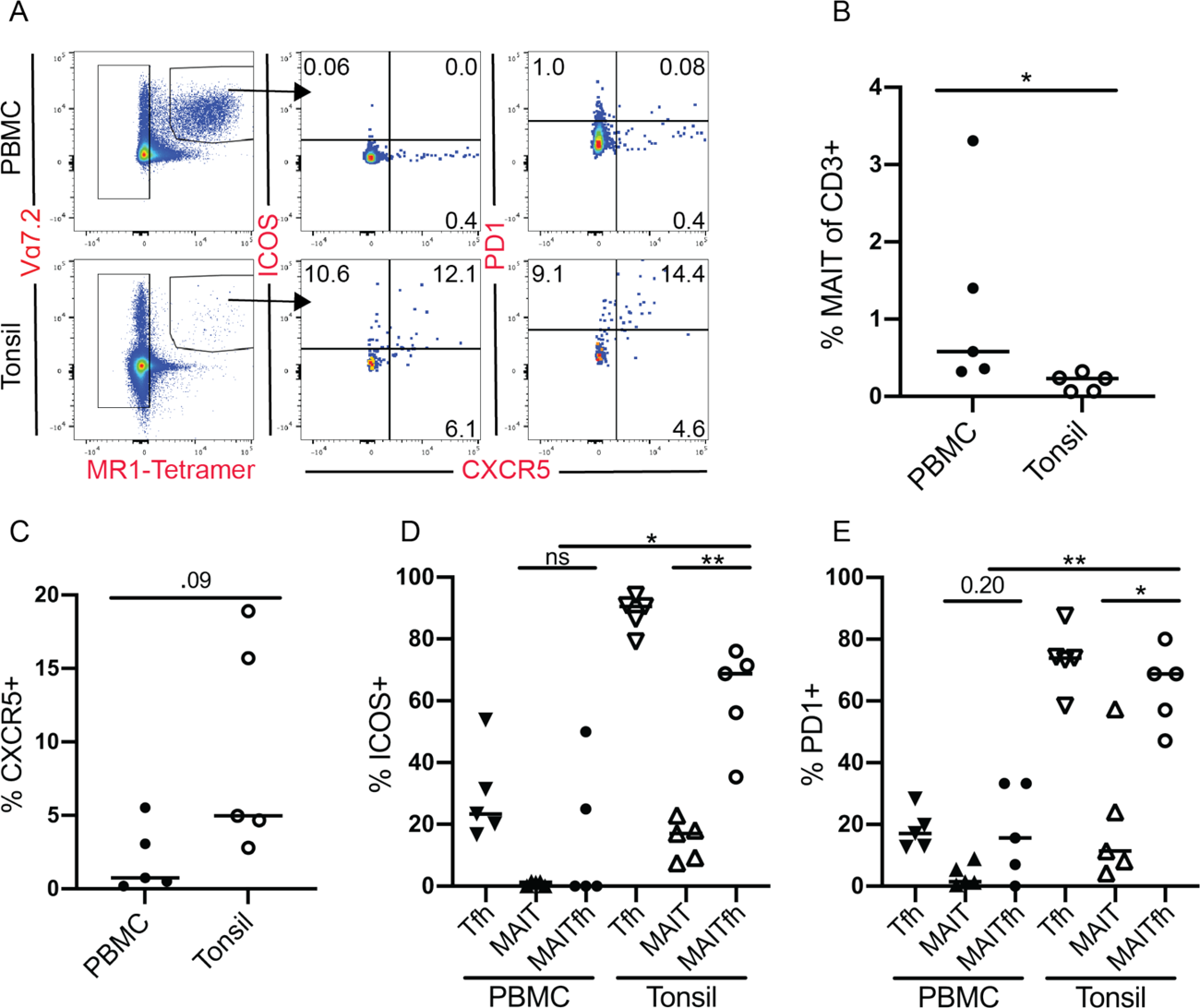
Tfh co-stimulatory marker expression in paired pediatric tonsil and PBMC samples. (**A**) Representative FACS plots of unstimulated paired pediatric CD3^+^ PBMC (top) and tonsil (bottom) donor cells with MAIT cell (gated in left panel) co-expression of CXCR5^+^ with ICOS (middle) and PD1 (right). (**B**) Quantification of MAIT cell frequency as percentage of CD3^+^. (**C**) Frequency of CXCR5 MAIT cells. Frequency of (**D**) ICOS and (**E**) PD1 MAIT, MAITfh and Tfh cells. Data are represented as Median from 2 pooled independent experiments. n *≥* 13. *\*p* < 0.05, *\*\*p*<0.01, *\*\*\*p*<0.001, *\*\*\*\*p*<0.0001 by two-tailed Mann-Whitney *U* test

**Supplementary Fig. 5.**
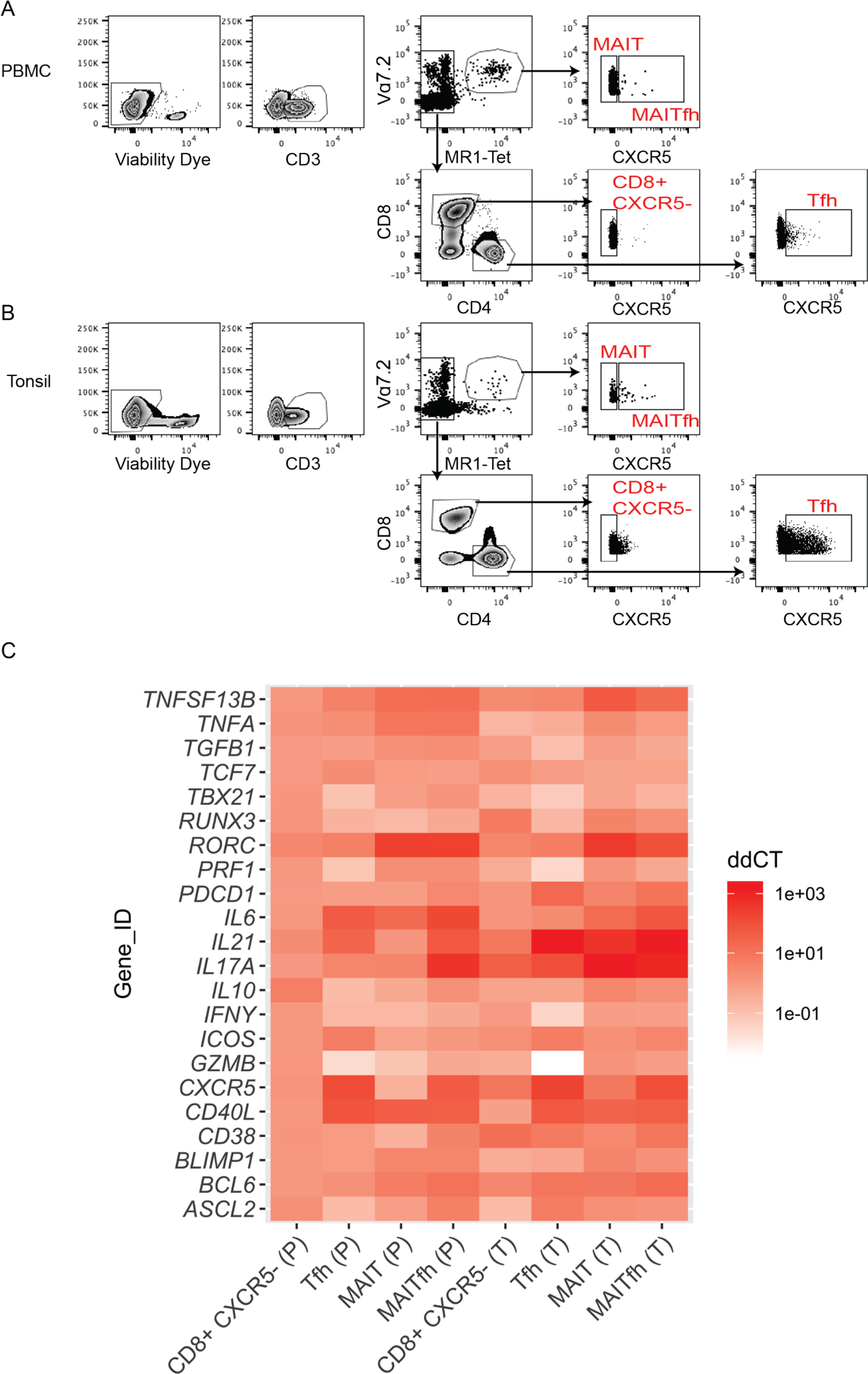
qPCR and flow gating strategy (Fig 2) and additional FAC sorted qPCR targets. Representative flow cytometry gating strategy for qPCR FAC sorted and flow populations in Fig.2 from (**A**) PBMC and (**B**) tonsils. (**C**) Heat map of additional qPCR targets described in Fig 2. As described, data are represented as *ΔΔ*Ct relative to *β*-Actin and then the PBMC-CD8^+^ CXCR5^-^ population. Rows denote gene ID and columns denote sample group (P=PBMC, T=tonsil). Heat map data are represented by mean of n=6-8 samples per group.

**Supplementary Fig. 6.**
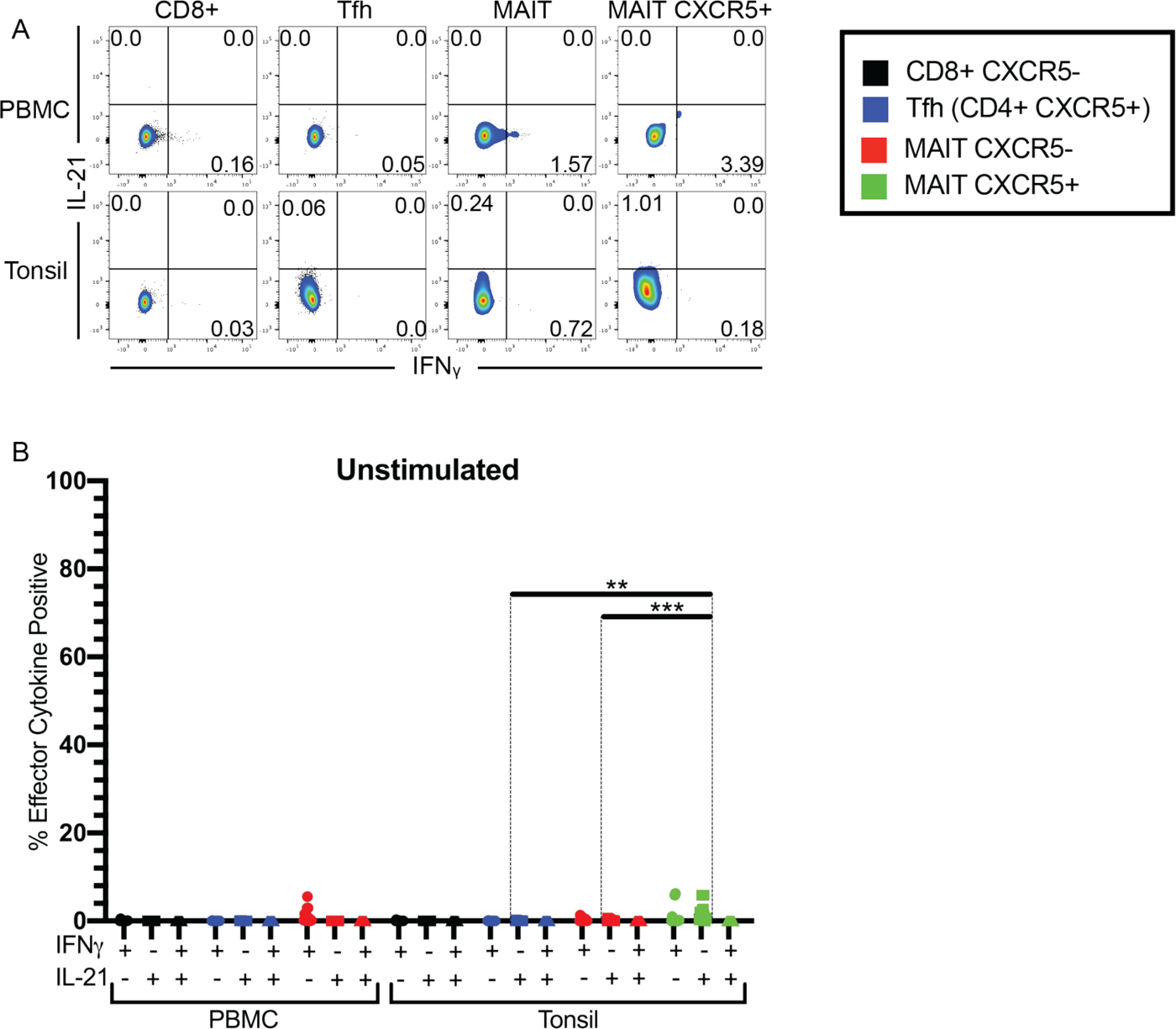
IFN*γ* and IL-21 expression in unstimulated PMBC and tonsil T cell populations. (**A**) Representative flow cytometry from 6 hour no stimulation controls with BFA added for final 4 hours in PBMC and tonsil T cell populations showing IL-21 and IFN*γ* co-expression. (**B**) Frequency of T cell populations expressing IL-21, IFN*γ* or both. Data are represented as median from two pooled independent experiments. n*≥*5. *\*p* < 0.05, *\*\*p*<0.01, *\*\*\*p*<0.001, *\*\*\*\*p*<0.0001 by two-tailed Mann-Whitney *U* test. Legend in black box denotes experimental groups.

**Supplementary Fig. 7.**
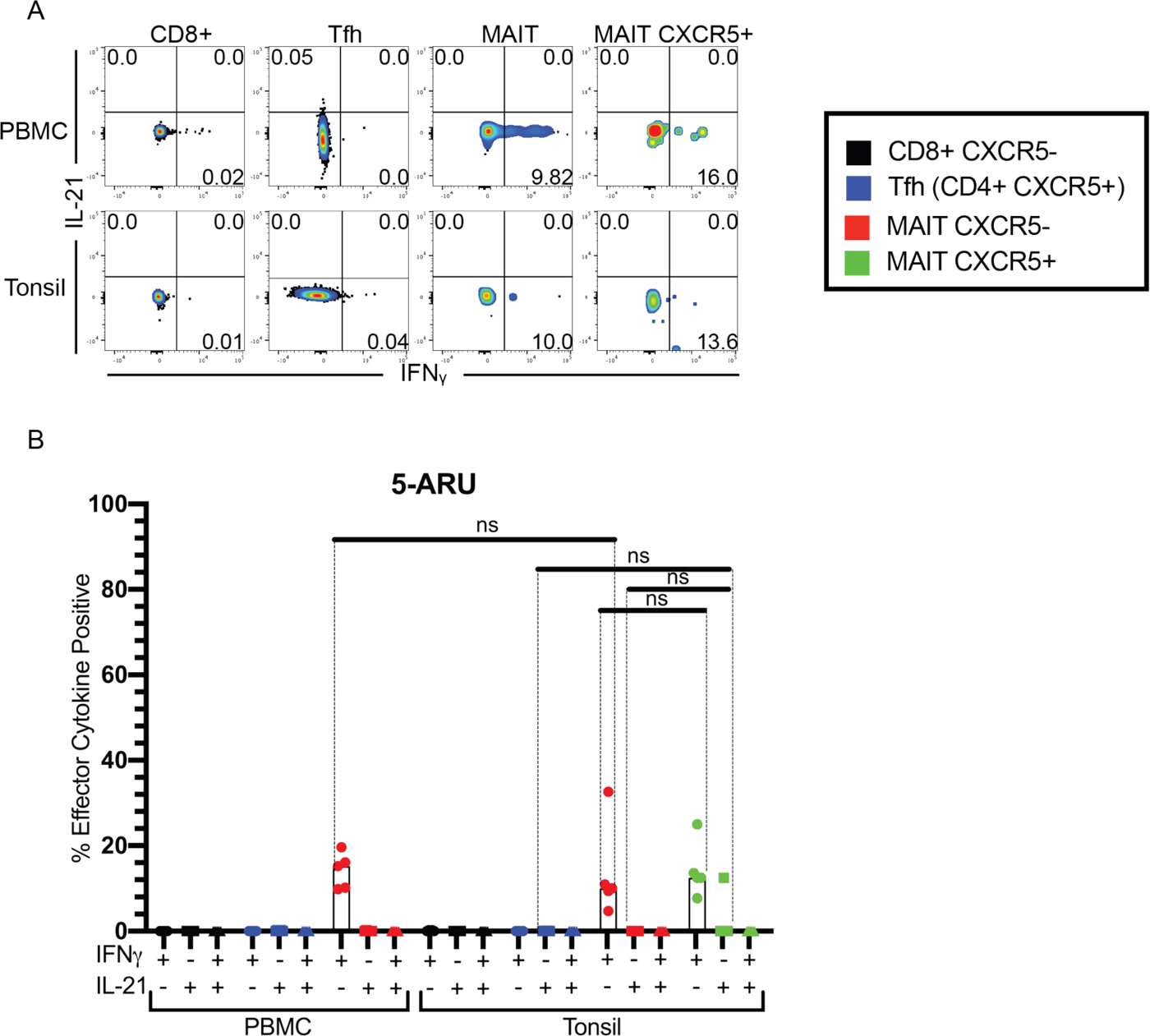
IFN*γ* and IL-21 expression in 5-A-RU stimulated PMBC and tonsil T cell populations. (**A**) Representative flow cytometry of 6 hr 5-A-RU (0.5 μM/ml) and Methylglyoxal (50 μg/ml) stimulated cultures with BFA in PBMC and tonsil T cell populations showing IL-21 and IFN*γ* co-expression. (**B**) Frequency of T cell populations expressing IL-21, IFN*γ*or both. Data are represented as median from one independent experiment. n=5. *\*p* < 0.05, *\*\*p*<0.01, *\*\*\*p*<0.001, *\*\*\*\*p*<0.0001 by two-tailed Mann-Whitney *U* test. Legend in black box denotes experimental groups.

**Supplementary Fig. 8.**
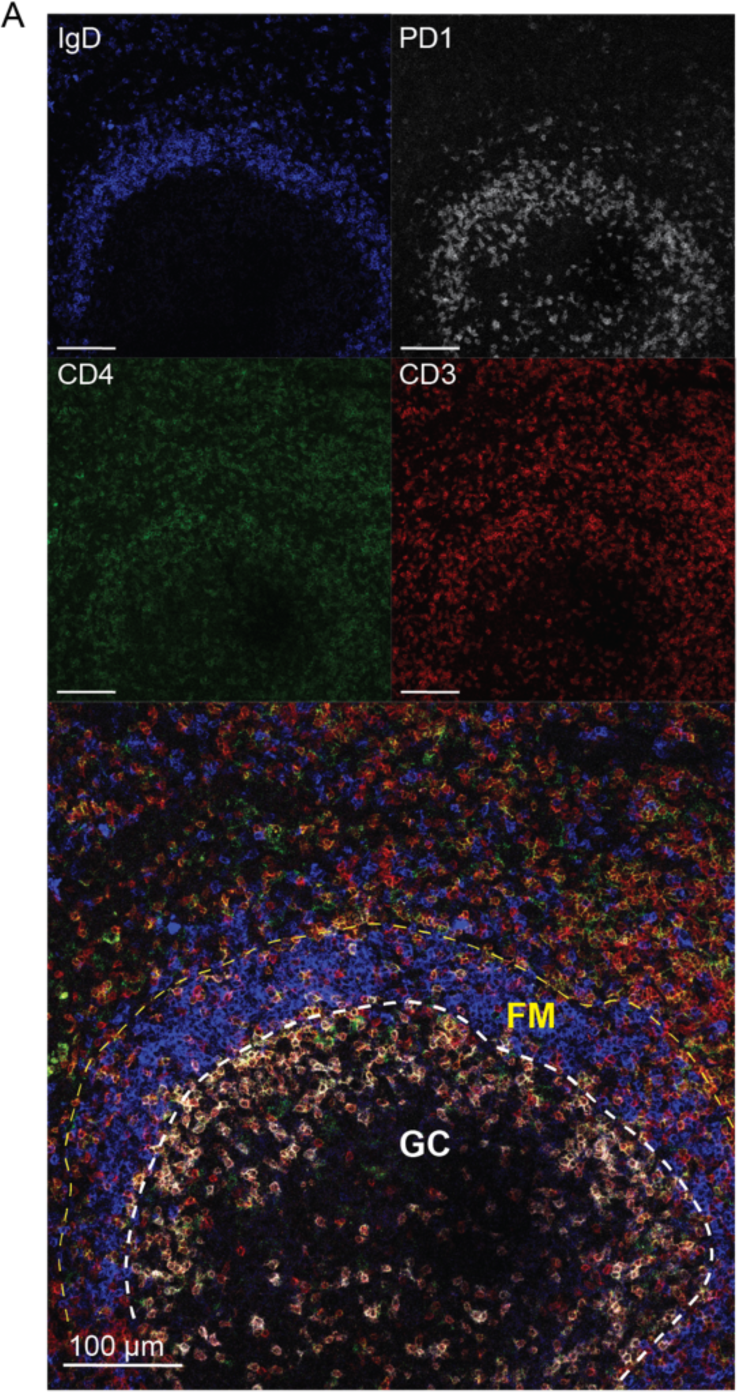
GC Tfh cells are marked by high PD1 expression in tonsils. (**A**) Representative immunofluorescence images from tonsil cryosections stained with anti-IgD (blue), anti-PD1 (white), anti-CD4 (green), and anti-CD3 (red) and imaged on a Leica SP8 confocal microscope with 20X objective. Enlarged composite image on bottom with a germinal center (GC) highlighted by PD1^High^ GC Tfh cells outlined with white dashed line, and the follicular mantle (FM) made up of IgD^+^ naïve B cells outlined with a yellow dashed line. All scale bars represent 100 μm.

**Supplementary Fig. 9.**
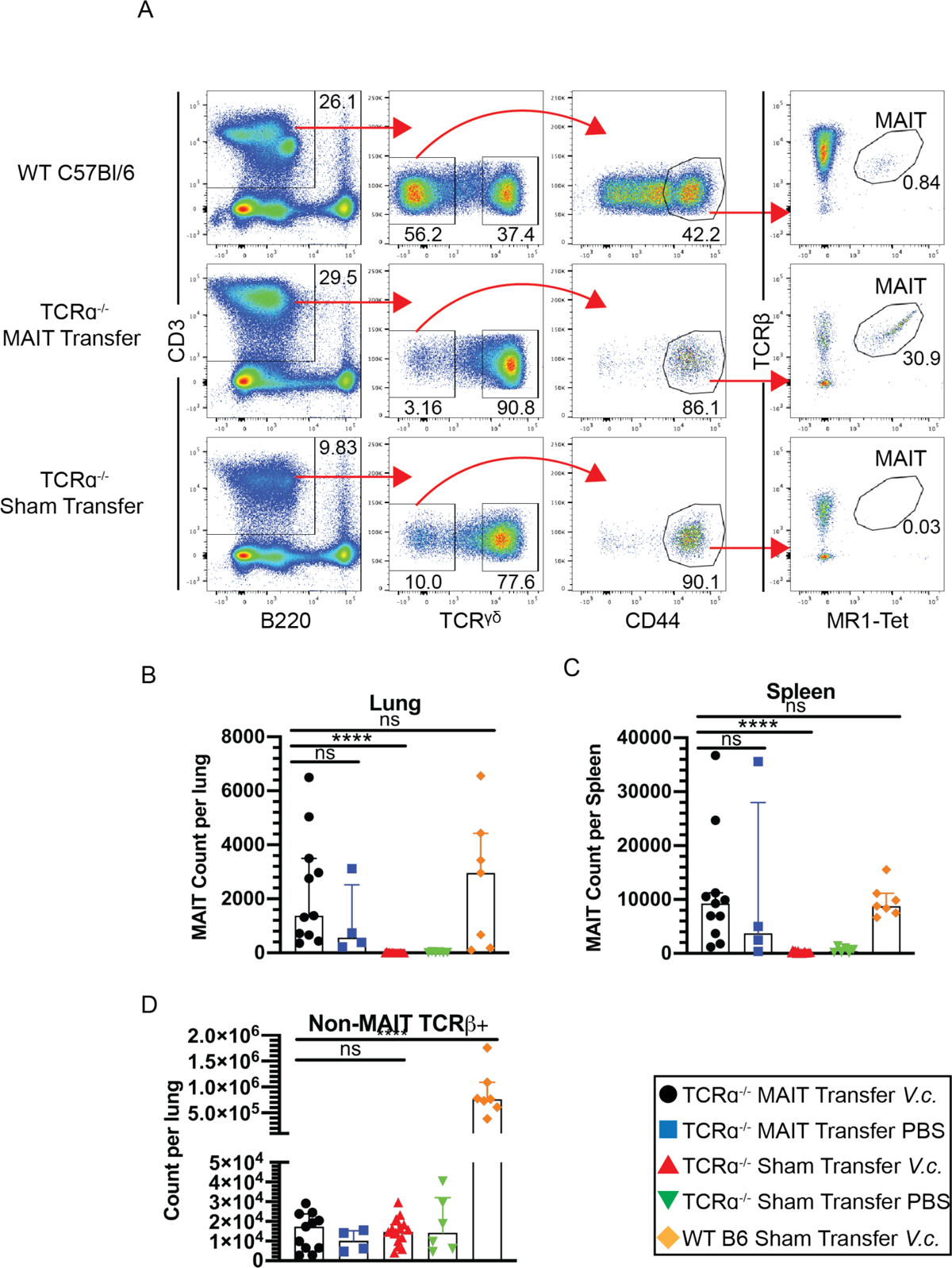
Flow gating for murine lung MAIT and T lymphocyte counts. (**A**) Representative flow gating of murine MAIT cells from digested lung tissue highlighted in Fig. 4a. MAIT cells were defined as live B220^-^ CD3^+^ TCR*γδ*^-^ CD44^High^ TCR*β*^+^ MR1-Tetramer^+^. Only WT-*V.c.*, MAIT-*V.c.*, and Sham-*V.c.* groups are portrayed. (**B**) Total MAIT cell count per lung. (**C**) Total MAIT cell count per spleen. (D) Total non-MAIT CD3^+^ TCR*β*^+^ T cells per lung. Data are represented as Median with IQR from 4 independent experiments. n= 4-15 mice per group. *\*p* < 0.05, *\*\*p*<0.01, *\*\*\*p*<0.001, *\*\*\*\*p*<0.0001 by two-tailed Mann-Whitney *U* test. Legend in black box denotes experimental groups.

**Supplementary Fig. 10.**
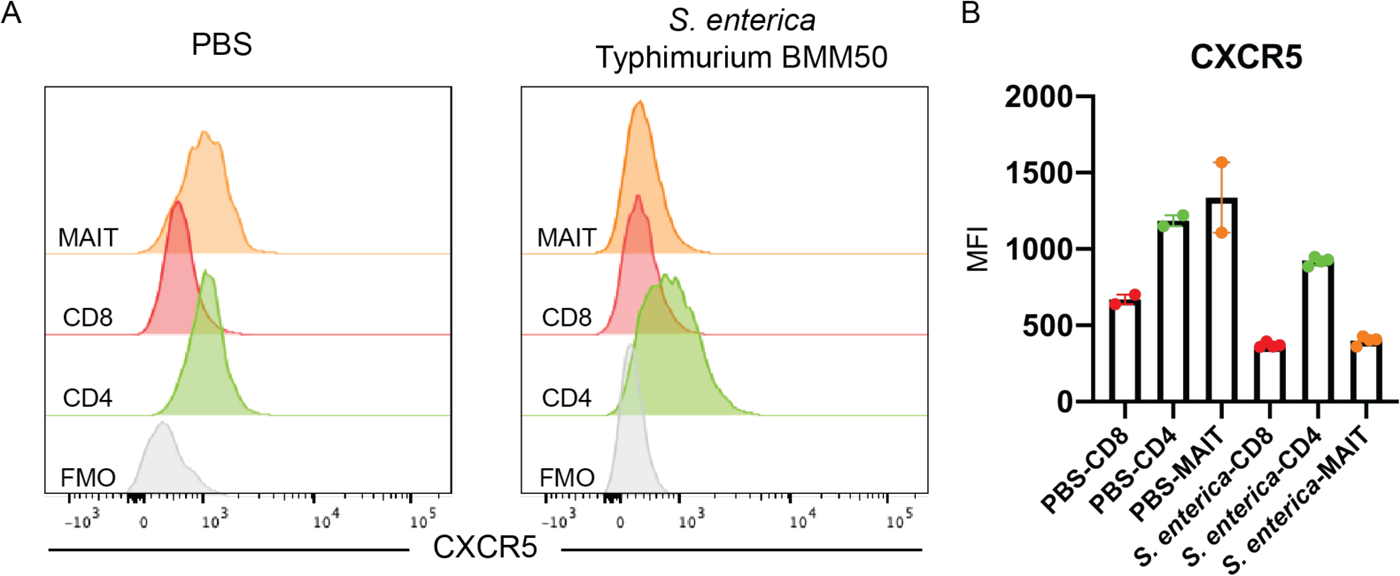
CXCR5 expression on *S. enterica* Typhimurium expanded MAIT cells prior to adoptive transfer. Representative flow cytometry histograms (**A**) and mean fluorescence intensity (**B**) of CXCR5 expression in T cell subsets of Day 7 WT C57Bl/6 mice given intranasal PBS (Left) or 10^6^ CFU *S.* Typhimurium (Left). Data are represented as Median from 1 experiment. n= 2-4 mice per group.

**Supplementary Fig. 11.**
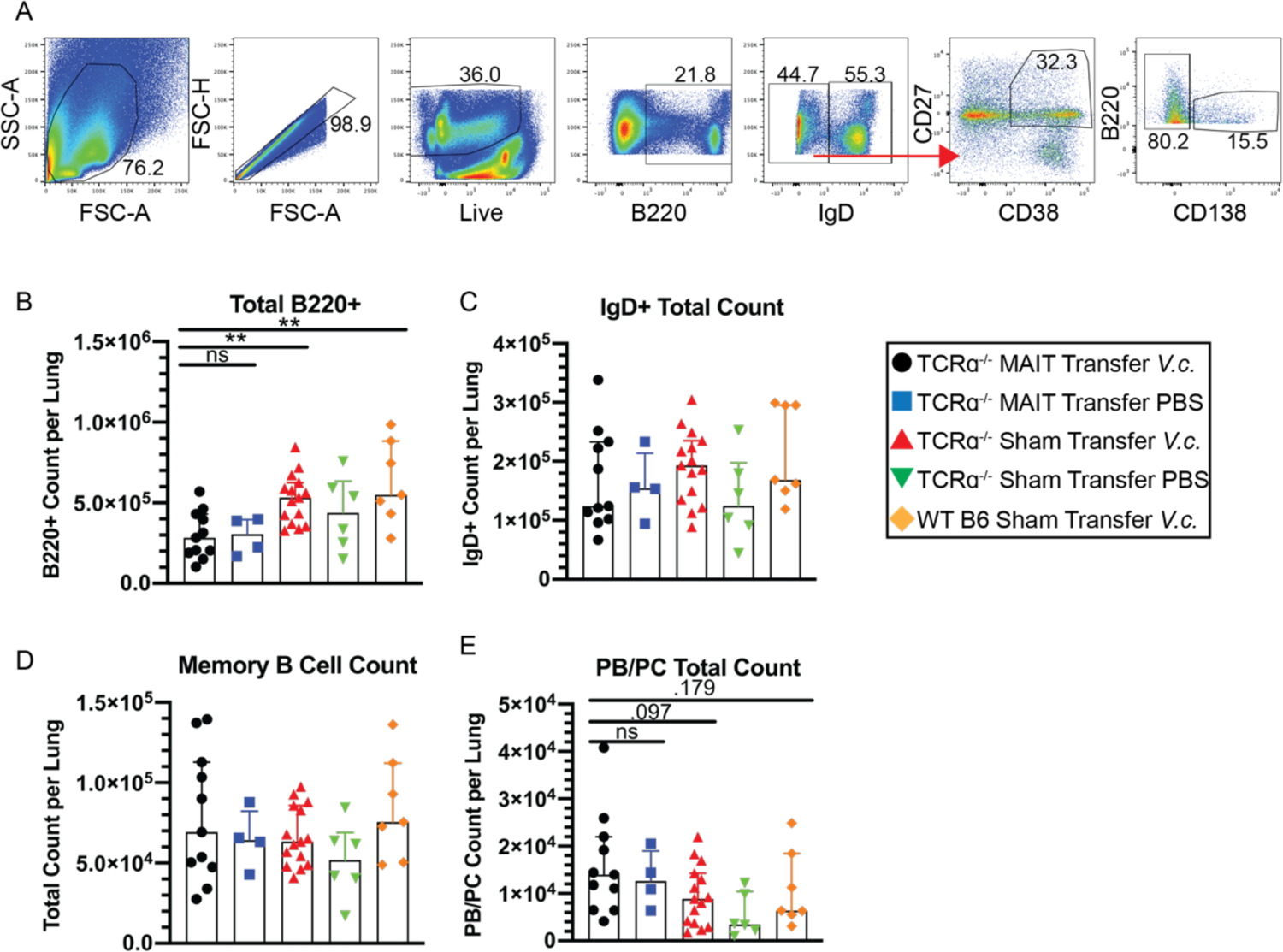
Flow gating for murine lung B lymphocyte counts. (**A**) Representative flow cytometry gating strategy of murine B cell subsets from digested lung tissue. (**B**) Total B220^+^ B cells per lung. (**C**) Total B220^+^ IgD^+^ Naïve B cells per lung. (**D**) Total B220^+^ IgD^-^ CD38^+^CD27^+^CD138^-^ Memory B cells per lung. (**E**) Total B220^low^ IgD^-^ CD38^+^CD27^+^CD138^+^ PB/PCs per lung. Data are represented as Median with IQR from 4 independent experiments. n= 4-15 mice per group. *\*p* < 0.05, *\*\*p*<0.01, *\*\*\*p*<0.001, *\*\*\*\*p*<0.0001 by two-tailed Mann-Whitney *U* test. Legend in black box denotes experimental groups.

**Supplementary Fig. 12.**
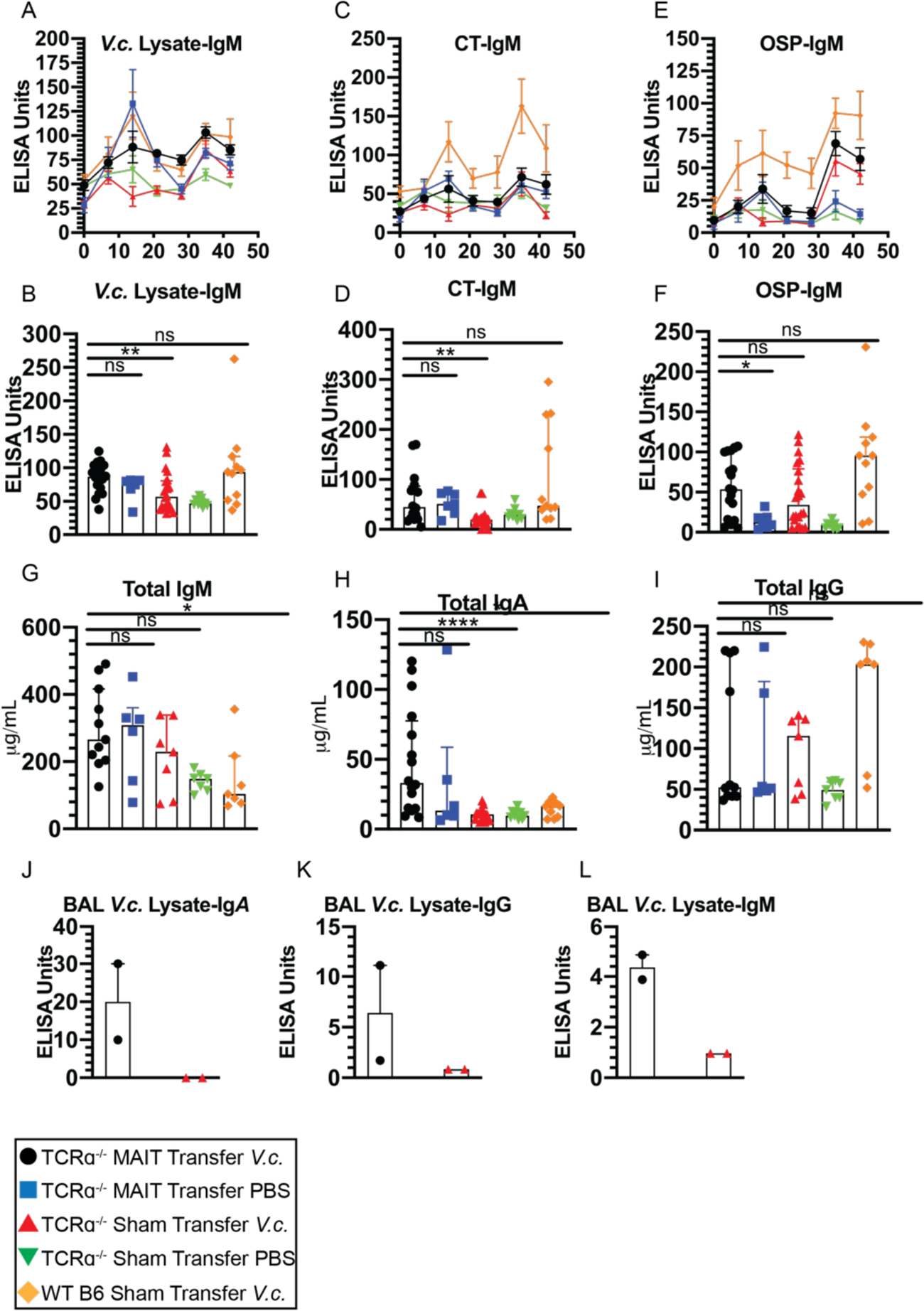
Serum IgM, total IgM, IgG and IgA and BAL fluid ELISAs. (A-F) Serum *V. cholerae* lysate, OSP, and CT IgM ELISAs representing weekly antibody kinetics (A, C & E) and Day 42 endpoint responses (B, D & F). Data are represented as ELISA units normalized to a pooled serum positive control from WT B6 mice challenged with V. cholerae. Total IgM (G), IgA (H) and IgG (I) ELISAs from Day 42 endpoint serum. Total Ig was quantitated using a standard curve based on known Ig quantities. Data are represented as Median with IQR from 5 independent experiments. n= 6-22 mice per group. *p < 0.05, **p<0.01, ***p<0.001, ****p<0.0001 by two-tailed Mann-Whitney U test. *V. cholerae* lysate IgA (J), IgG (K) and IgM (L) ELISAs from D42 BAL fluid. Legend in black box denotes experimental groups.

